# Alterations in the amplitude and burst distribution of sensorimotor beta oscillations impair reward-dependent motor learning in anxiety

**DOI:** 10.1101/442772

**Authors:** Sebastian Sporn, Thomas P. Hein, María Herrojo Ruiz

## Abstract

Anxiety results in sub-optimal motor performance and learning; yet, the precise mechanisms through which these modifications occur remain unknown. Using a reward-based motor sequence learning paradigm, we show that concurrent and prior anxiety states impair learning by biasing estimates about the hidden performance goal and the stability of such estimates over time (volatility). In an electroencephalography study, three groups of participants completed our motor task, which had separate phases for motor exploration (baseline) and reward-based learning. Anxiety was manipulated either during the initial baseline exploration phase or while learning. We show that anxiety induced at baseline reduced motor variability, undermining subsequent reward-based learning. Mechanistically, however, the most direct consequence of state anxiety was an underestimation of the hidden performance goal and a higher tendency to believe that the goal was unstable over time. Further, anxiety decreased uncertainty about volatility, which attenuated the update of beliefs about this quantity. Changes in the amplitude and burst distribution of sensorimotor and prefrontal beta oscillations were observed at baseline, which were primarily explained by the anxiety induction. These changes extended to the subsequent learning phase, where phasic increases in beta power and in the rate of long (> 500 ms) oscillation bursts following reward feedback were linked to smaller updates in predictions about volatility, with a higher anxiety-related increase explaining the biased volatility estimates. These data suggest that state anxiety alters the dynamics of beta oscillations during general performance, yet more prominently during reward processing, thereby impairing proper updating of motor predictions when learning in unstable environments.

## Introduction

Anxiety involves anticipatory changes in physiological and psychological responses to an uncertain future threat (Bishop, 2007; Grupe and Nitschke, 2013). Previous studies established that trait anxiety interferes with prefrontal control of attention in perceptual tasks, whereas state anxiety modulates the amygdala during detection of threat-related stimuli (Bishop, 2007; Bishop, 2009). Computational modeling work has started to examine the mechanisms through which anxiety might impair learning, revealing that highly anxious individuals do not correctly estimate the degree of uncertainty in the environment during aversive learning (Browning et al. 2015). In the area of motor control, research has shown that stress and anxiety have detrimental effects on performance (Baumeister, 1984; Beilock and Carr, 2001). These results have been interpreted as the interference of anxiety with information-processing resources; also as a shift towards an inward focus of attention and an increase in conscious processing of the movement (Eysenck & Calvo, 1992; Pijpers et al., 2005). The effects of anxiety on motor learning, however, are often inconsistent, and a mechanistic understanding is still lacking. Delineating mechanisms through which anxiety influences motor learning is important to ameliorate its impact in different settings, including in motor rehabilitation programs.

Motor variability could be one component of motor learning that is affected by anxiety; it is defined as the variation of performance across repetitions (van Beers et al., 2004), and is affected by various factors including sensory and neuromuscular noise (He et al., 2016). As a form of action exploration, movement variability is increasingly recognized to benefit motor learning (Todorov and Jordan, 2002; Wu et al., 2014; Pekny et al., 2015), particularly during reward-based learning, with discrepant effects in motor adaptation paradigms (He et al., 2016; Singh et al., 2016). These findings are consistent with the vast amount of research on reinforcement learning, demonstrating increased learning following initial exploration (Sutton and Barto, 1998).

Yet contextual factors can reduce variability. For instance, state anxiety leads to ritualistic behavior, characterized by movement redundancy, repetition, and rigidity (Lang et al., 2015). This finding resembles the reduction in behavioral variability and exploration that manifests across animal species in stressful environments (Morgan and Tromborg, 2007). Based on these results, we set out to test the hypothesis that state anxiety modulates motor learning through a reduction in motor variability.

A second component that could be influenced by anxiety is the flexibility to adapt to changes in the task structure during learning. Individuals affected by anxiety disorders exhibit an intolerance of uncertainty, which contributes to excessive worry and emotional dysregulation (Ouellet et al., 2019). Turning to non-clinical populations, computational studies established that highly anxious individuals exhibit difficulties in estimating environmental uncertainty both in aversive and reward-based tasks (Browning et al., 2015; Huang et al., 2017). Failure to adapt to volatile or unstable environments thus impairs learning of action-outcome contingencies in these settings. Accordingly, in the context of motor learning, and more specifically, reward-based motor learning, we proposed that an increase in anxiety would affect individuals’ estimation of uncertainty about the stability of the task structure, such as the rewarded movement.

On the neural level, we posited that changes in motor variability are driven by neural variability in premotor and motor areas. Support for our hypothesis comes from animal studies demonstrating that variability in the primate premotor cortex tracks behavioral variability during motor planning (Churchland et al., 2006). Further evidence supports that changes in variability in single-neuron activity in motor cortex drive motor exploration during initial learning, and reduce it following intensive training (Mandelblat-Cerf et al., 2009; Santos et al., 2015). Additionally, the basal ganglia are crucial for modulating variability during learning and production, as shown in songbirds and, indirectly, in patients with Parkinson’s disease (Kao et al., 2005; Ölveczky et al., 2005; Pekny et al., 2015).

In the present study, we analyzed sensorimotor beta oscillations (13-30Hz) as a candidate mechanism driving motor exploration and variability. Beta oscillations modulate different aspects of performance and motor learning (Herrojo Ruiz et al., 2014; Bartolo and Merchant, 2015; Tan et al., 2014), as well as reward-based learning (Haji Hosseini et al., 2012). Increases in beta power following movement have been proposed to signal higher reliance on prior information about the optimal movement (Tan et al., 2016), which would reduce the impact of new evidence on the update of motor commands. We therefore tested the additional hypothesis that changes in beta oscillations mediate the effect of anxiety on belief updates and the estimation of uncertainty driving reward-based motor learning. Although power changes were traditionally the primary focus of research on oscillations, there is a renewed interest towards assessing dynamic properties of oscillatory activity, such as the presence of brief bursts (Poil et al., 2008). Brief oscillation bursts are considered to be a central feature of physiological beta in motor-premotor cortex and the basal ganglia (Feingold et al., 2015; Tinkhauser et al., 2017; Little et al., 2018). The assessment of power and burst distribution of beta oscillations thus allows us to capture dynamic changes in neural activity induced by anxiety and their link to behavioral effects.

To test our hypotheses, we recorded electroencephalography (EEG) in three groups of participants while they completed a reward-based motor sequence learning paradigm, with separate phases for motor exploration (baseline) and reward-based learning. We manipulated anxiety by informing participants about an upcoming public speaking task (Lang et al., 2015). Using a between-subject design, the anxiety manipulation targeted either the baseline or the reward-based learning phase. Analysis of the EEG signals aimed to assess anxiety-related changes in the power and burst distribution in beta oscillations in relation to changes in behavioral variability and reward-based learning.

## Results

Sixty participants completed our reward-based motor sequence learning task, consisting of three blocks of 100 trials each over two phases (**Figure 1**): a baseline motor exploration (block 1) and a reward-based learning phase (blocks 2 and 3: termed training thereafter). Prior to the experimental task, we recorded in each participant 3 min of EEG at rest with eyes open. Next, on a digital piano, participants played two different sequences of seven and eight notes during the exploration and training phases respectively (**Figure 1A**). The sequence patterns were designed so that the key presses would span a range of four neighboring keys on the piano. Participants were explicitly taught the tone sequences prior to the start of the experiment, yet precise instructions about the timing or loudness (keystroke velocity, Kvel) were not provided. The rationale for selecting two different sequences for the baseline and training phases was to avoid carry-over effects of learning or a preferred performance pattern from the baseline period into the reward-based learning phase (following Wu et al., 2014).

**Figure 1.**
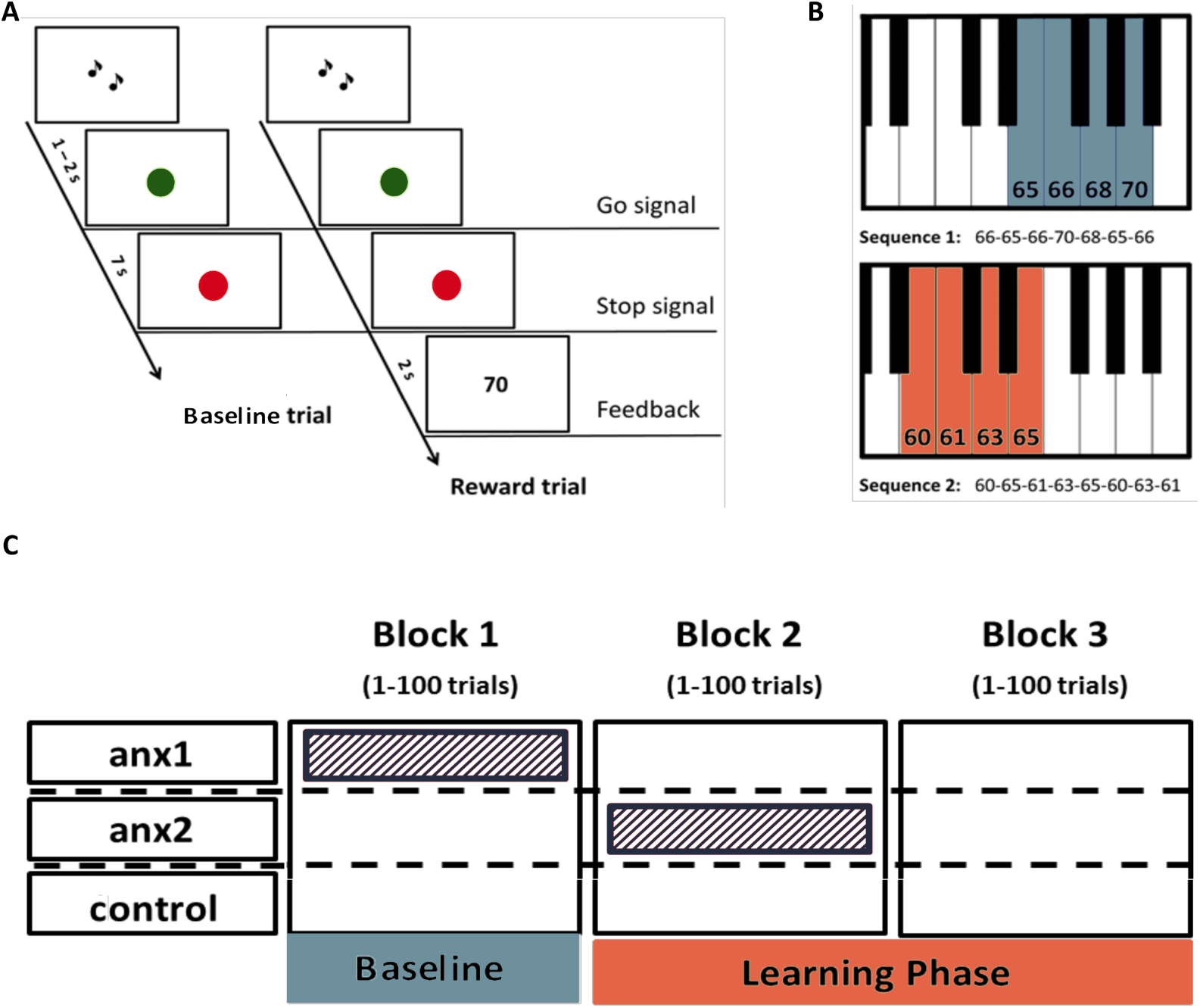
A Novel Paradigm for Testing Reward-Based Motor Sequence Learning. (A) Schematic of the task. Participants played sequence1 during 100 baseline exploration trials, followed by 200 trials over two blocks of reward-based learning performing sequence2. During the learning blocks, participants received a performance-related score between 0-100 that would lead to monetary reward. (B) Pitch content of the sequences used in the baseline exploration (sequence1) and reward-based learning blocks (sequence2), respectively. (C) Schematic of the anxiety manipulation. The shaded area denotes the phase in which anxiety was induced in each group, using the threat of an upcoming public speaking task, which took place immediately after that block was completed.

During the baseline exploration phase, participants were informed they could freely change the pattern of temporal intervals between key presses (inter-keystroke intervals, IKIs) and/or the loudness of the performance every trial, and that no reward or feedback would be provided. During training performance-based feedback in the form of a 0-100 score was provided at the end of each trial. Participants were informed that the overall average score would be translated into monetary reward. They were directly instructed to explore the temporal or loudness dimension (or both) and to use feedback scores to discover the unknown performance objective (which, unbeknownst to them, was related to the pattern of IKIs). The task-related dimension was therefore timing (**Figure 2**), whereas keystroke velocity was the non-task related dimension.

**Figure 2.**
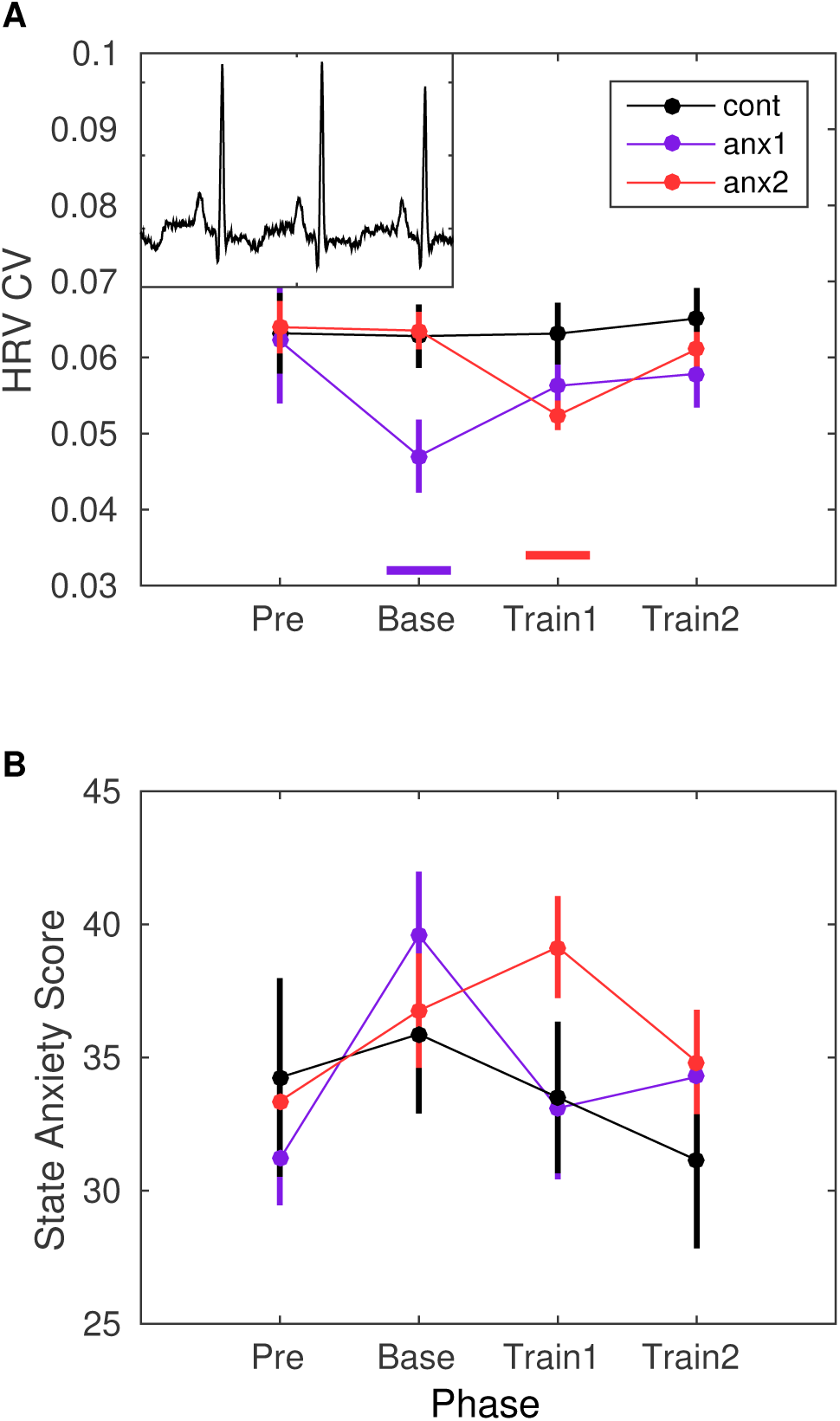
Heart-rate varibility (HRV) modulation by the anxiety manipulation. (A) Average HRV measured as the coefficient of variation of the inter-beat-interval is displayed across the experimental blocks: initial resting state recording (Pre), baseline exploration (Base), first block of training (Train1) and, last block of training (Train2). Relative to Pre, there was a significant drop in HRV in anx1 participants during baseline exploration (P_FDR_ < 0.05, Δ_dep_ = 0.81, CI = [0.75, 0.87]). In anx2 participants the drop in HRV was found during the first training phase, which was targeted by the anxiety manipulation (P_FDR_ < 0.05, FDR-corrected, Δ_dep_ = 0.78, CI = [0.71, 0.85]). Between-group comparisons revealed that anx1, relative to the control group, exhibited a significantly lower HRV during baseline exploration (P_FDR_ < 0.05, Δ = 0.75, CI = [0.65, 0.85], purple bar at the bottom). The anx2 group manifested a significant drop in HRV relative to controls during the first training block (P_FDR_ < 0.05, Δ = 0.71, CI = [0.62, 0.80], red bar at the bottom). These results demonstrate a group-specific modulation of anxiety relative to controls during the targeted blocks. The mean HR did not change within or between groups (P > 0.05). **(B)** STAI state anxiety score in each group across the different experimental phases. Participants completed the STAI state anxiety subscale first at the start of the experiment before the resting state recording (Pre) and subsequently again immediately before each experimental block (and right after the anxiety induction: Base, Train1, Train2). There was a within-group significant increase in the score for each experimental group during the phase targeted by the anxiety manipulation (anx1: Bas relative to Pre, average score 40[2] and 31[2], respectively; P_FDR_ < 0.05, Δ_dep_ = 0.74, CI = [0.68, 0.80]; anx2: Train1 relative to Pre, average score 39[2] and 34[2], respectively; P_FDR_ < 0.05, Δ_dep_ = 0.78, CI = [0.68, 0.86]). Between-group differences were non-significant.

The performance measure that was rewarded during training was the vector norm of the pattern of temporal differences between adjacent IKIs (See *Materials and Experimental design).* Similar combinations of IKIs could lead to the same rewarded norm of IKI-difference values, and therefore to the same score. Participants were unaware of the existence of these multiple solutions. The multiplicity in the mapping between performance and score could lead participants to perceive an increased level of volatility in the environment (changes in the rewarded performance over time). This motivated us to assess their estimation of volatility during reward-based learning and its modulation by anxiety. In addition, we investigated whether higher baseline variability would lead to higher scores during subsequent reward-based learning, independently of changes in variability during this latter phase. If initial baseline exploration improves learning of the mapping between the actions and their sensory consequences, then participants could learn better from performance-related feedback during the training phase regardless of their use of variability in this phase. Alternatively, it could be that participants who also use more variability during training discover the hidden goal by chance.

Participants were pseudo-randomly allocated to either a control group or to one of two experimental groups (**Figure 1B**): anxiety during exploration (anx1); anxiety during the first block of training (anx2). We measured changes in heart-rate variability (HRV) and heart-rate (HR) four times throughout the experimental session: resting state (3 min, prior to performance blocks); block1; block2; block3. In addition, the state subscale from the State-Trait Anxiety Inventory (STAI, state scale X1, 20 items; Spielberger, 1970) was assessed four times: prior to the resting state recording and also immediately before the beginning of each block, and thus after the induction of anxiety in the experimental groups. The HRV index and STAI state anxiety subscale were able to dissociate in each experimental group the phase targeted by the anxiety manipulation and the initial resting phase (within-group effects, **Figure 2A-B**). These results confirmed that the experimental manipulation succeeded in inducing physiological and psychological responses within each experimental group consistent with an anxious state during the targeted phase, as reported previously (Feldman et al., 2004).

Statistical analysis of behavioral and neural measures focused on the separate comparison between each experimental group and the control group (contrasts: anx1 – controls, anx2 – controls). See **Methods and Materials.**

### Behavioral Results

#### Lower baseline task-related variability is associated with poorer reward-based learning

All groups of participants demonstrated significant improvement in the achieved scores during reward-based learning, confirming they effectively used feedback to approach the hidden target performance (changes in average score from block 2 to block3; anx1: *P* = 0.008, non-parametric effect size estimator for dependent samples, Δ_dep_ 0.93, confidence interval or CI = [0.86, 0.99]; anx2: *P* = 0.004, Δ_dep_= 0.83, CI = [0.61, 0.95]; controls: *P* = 0.001, Δ_dep_= 0.92, CI = [0.72, 0.98]).

Assessment of motor variability was performed separately in the task-related temporal dimension and the non-task-related keystroke velocity dimension. Temporal variability – and similarly for keystroke velocity – was estimated using the across-trials coefficient of variation of IKI (termed cvIKI thereafter; **Figure 3A-B**). This index was computed in bins of 25 trials, which therefore provided four values per experimental block. We hypothesized that in the total population a higher degree of task-related variability at baseline (that is, playing different temporal patterns in each trial), and therefore higher cvIKI, would improve subsequent reward-based learning, as this latter phase rewarded the temporal dimension. A non-parametric rank correlation analysis across the 60 participants revealed that participants who achieved higher scores in the training phase exhibited a larger across-trials cvIKI at baseline (Spearman ρ = 0.45, *P* = 0.003; **Figure 3C**). A similar result was obtained when excluding anx1 participants from the correlation analysis, supporting that in the subsample of 40 participants who did not undergo the anxiety manipulation at baseline there was a significant association between the level of task-related variability and the subsequent score (ρ = 0.41, *P* = 0.04). No significant rank correlation was found between the scores and cvKvel.

**Figure 3.**
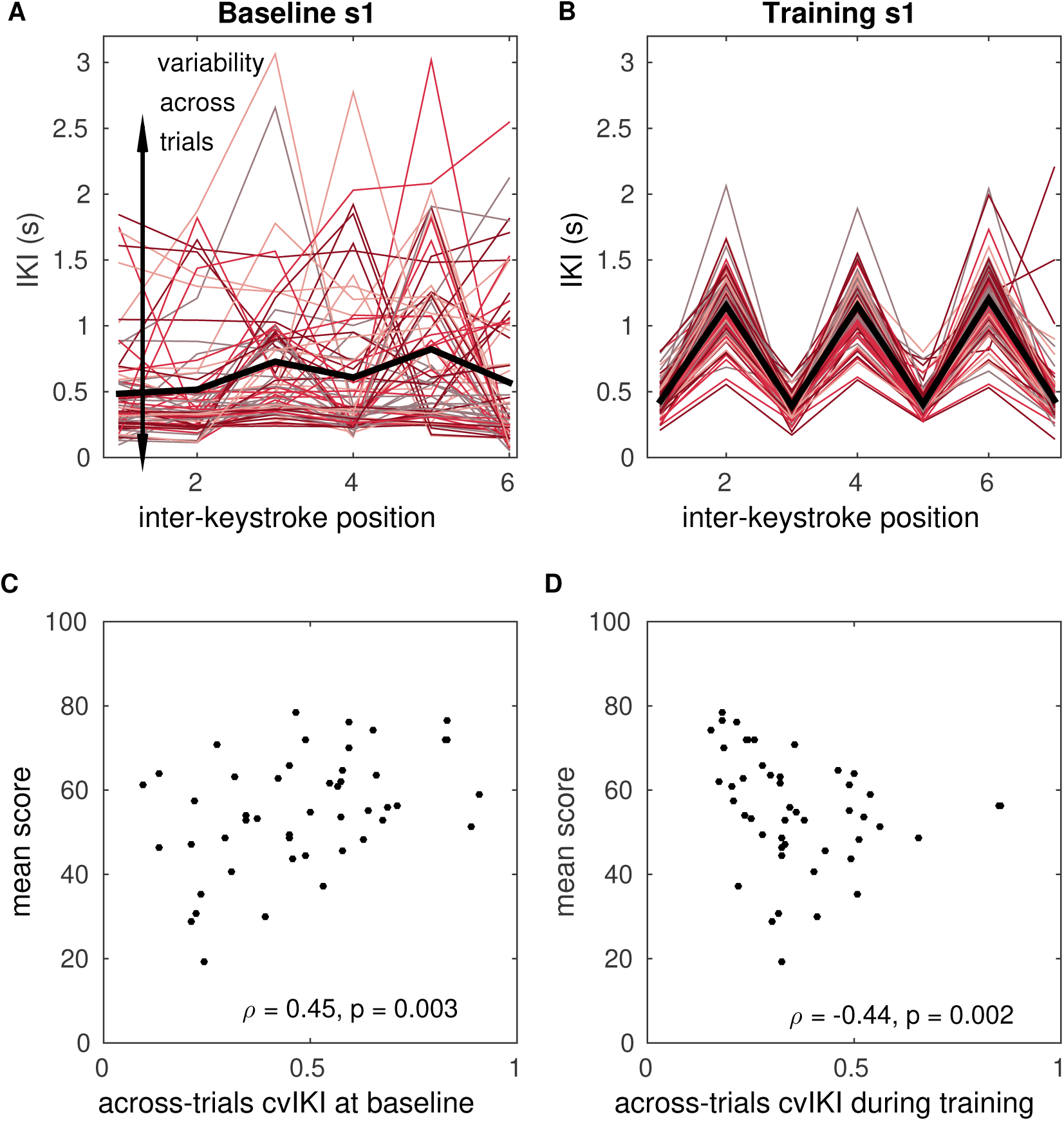
Temporal variability at baseline and during reward-based learning. (A-B) Illustration of timing performance during baseline exploration (A) and training (B) blocks in one representative participant, s1. The x-axis represents the position of the inter-keystroke interval (sequence1: 7 notes, corresponding to 6 inter-keystroke temporal intervals; sequence2: 8 notes, 7 inter-keystroke intervals). The y-axis shows the inter-keystroke interval (IKI) in ms. Black lines represent the mean IKI pattern. Task-related temporal variability was measured using the across-trials coefficient of variation of IKI, cvIKI. **(C)** Non-parametric rank correlation in the total population (N = 60) between the across-trials cvIKI at baseline and the average score achieved subsequently during training (Spearman ρ = 0.45, p = 0.003). **(D)** Same as C but using the individual value of the across-trials cvIKI from the training phase (Spearman ρ = −0.44, p = 0.002). Bars around the mean display ±SEM.

We also assessed whether the degree of cvIKI during training was associated with the average score and found an inverted pattern: There was a significant negative non-parametric rank correlation between the cvIKI index and the mean score (ρ= −0.44, *P* = 0.002; **Figure 3D**). A significant effect was not found for the cvKvel parameter (*P* > 0.05).

Notably, the amount of variability in timing and keystroke velocity used by participants was not correlated (cvIKI and cvKvel: ρ = 0.021, *P* = 0.788). This indicates that in our task participants could vary the temporal and velocity dimensions separately. Finally, the degree of cvIKI during training and baseline were not correlated (ρ = 0.029, *P* = 0.848). These findings support that achieving higher scores during reward-based learning in our paradigm cannot be accounted for by a general tendency to explore more throughout all experimental blocks. In fact, larger variability during training was detrimental to discover and maintain the performance close to the target (**Figure 3D**).

### Anxiety at baseline reduces task-related variability and impairs subsequent reward-based learning

We assessed pair-wise differences in the behavioral measures between the control group and each experimental group (anx1, anx2), separately. Participants affected by state anxiety at baseline (anx1) achieved significantly lower scores in the subsequent reward-based learning phase relative to control participants (**Figure 4A**: *P_FDR_* < 0.05, Δ = 0.78, CI = [0.54, 0.92]). By contrast, in the anx2 group scores did not statistically differ from the scores in the control group (*P_FDR_* > 0.05). A planned comparison between both experimental groups demonstrated significantly higher scores in anx2 than in anx1 (*P_FDR_* < 0.05, Δ= 0.67, CI = [0.51, 0.80]).

**Figure 4.**
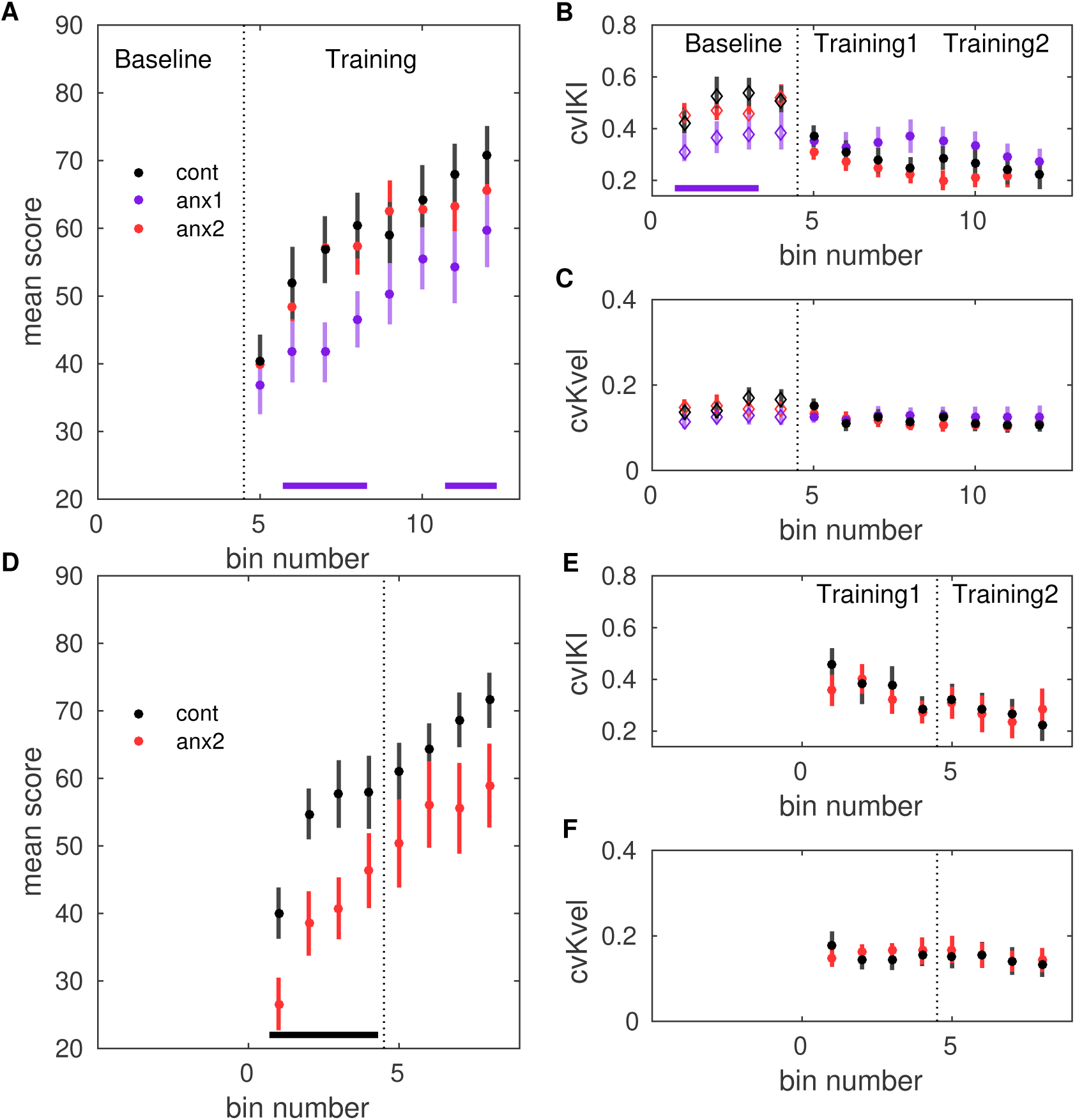
Effects of anxiety on behavioral variability and reward-based learning. The score was computed as a 0-100 normalized measure of proximity between the norm of the pattern of *differences* in inter-keystroke intervals performed in each trial and the target norm. **(A)** Scores achieved by participants in the anx1, anx2, and control groups across bins 5:12 (bins of 25 trials: trial range 101-300), corresponding to blocks 2 and 3 and the training phase. Participants in anx1 achieved significantly lower scores than control participants (P_FDR_ < 0.05, denoted by the bottom purple line). **(B)** Changes in across-trials cvKvel, revealing a signifcant drop in task-related exploration at baseline in anx1 relative to control participants (P_FDR_ < 0.05). Anx2 participants did not differ from control participants. **(C)** Same as (B) but for the across-trials cvKvel. **(D-F)** Control experiment: Effect of anxiety on variability and learning after removal of the baseline exploration phase. Panels D-F are displayed as panels A-C. Significant between-group differences are denoted by the black bar at the bottom (P_FDR_ < 0.05, Δ = 0.71, CI = [0.64, 0.78]). (F) In anx2 participants there was a significant drop in the mean scores during the first training block relative to control participants (P_FDR_ < 0.05, Δ = 0.77, CI = [0.68, 0.86]). Bars around the mean show ±SEM.

At baseline, anx1 used a lower degree of cvIKI than the control group (**Figure 4B**; *P_FDR_* < 0.05; Δ = 0.67, CI = [0.52, 0.85]). There was no between-groups (anx1, controls) difference in cvKvel (**Figure 4C**; *P_FDR_ >* 0.05). Performance at baseline in anx2 did not significantly differ from performance in the control group, both for cvIKI or cvKvel (*P_FDR_>* 0.05).

During the training blocks, there were no significant between-group differences in cvIKI or cvKvel (*P*_FDR_ > 0.05). There was, however, a significant and very pronounced drop in the use of temporal variability from the baseline to the training phase in control and anx2 participants (large effect sizes: *P* = 0.0078, Δ_dep_ = 0.81, CI = [0.62, 0.95], in controls; *P* = 0.0026, Δ_dep_ = 0.83, CI = [0.61, 0.90], in anx2). This drop corresponded to a change from a largely explorative regime at baseline (characterized by higher cvIKI) to a more constrained explorative regime early during training, followed by a gradual transition to the exploitation of the rewarded options in these groups (significant drop in cvIKI from block2 to block 3 in control and anx2 participants, respectively; *P* = 0.04, Δ_dep_ = 0.77, CI = [0.53, 0.87], in controls; *P* = 0.0054, Δ_dep_ = 0.83, CI = [0.62, 0.94], in anx2). In anx1 participants, by contrast, there was no significant change in cvIKI from the baseline to the training phase, or from block 2 to block3 during training. This outcome indicated that anx1 participants did not adapt their use of temporal variability to the task requirements.

Detailed analyses of the trial-by-trial changes in scores and performance using a Bayesian learning model and their modulation by anxiety are reported below.

General performance parameters, such as the average performance tempo or the mean keystroke velocity did not differ between groups, either during baseline exploration or training (*P* > 0.05). Participants completed sequence1 on average in 3.0 (0.1) seconds, between 0.68 (0.05) and 3.68 (0.10) s after the GO signal (non-significant differences between groups, *P* > 0.05). During training, they played sequence2 with an average duration of 4.7 (0.1) s, between 0.72 (0.03) and 5.35 (0.10) s (non-significant differences between groups, *P* > 0.05). The *mean* learned solution in each group was not significantly different, either during the first or second training block (*P* > 0.05; **Figure 4 – figure supplement 1**; but see trial-by-trial changes below).

These outcomes demonstrate that in our paradigm state anxiety reduced task-related motor variability when induced at baseline and this effect was associated with lower scores during subsequent reward-based learning. State anxiety, however, did not modulate task-related motor variability or the achieved scores when induced during reward-based learning. Finally, the different experimental manipulations did not affect the mean learned solution in each group.

### State anxiety during reward-based learning reduces learning rates if there is no prior baseline exploration

Because anx2 participants performed at a level not significantly different from that found in control participants during training, we asked whether the initial unconstrained motor exploration at baseline might have counteracted the effect of anxiety during training. Alternatively, it could be that the anxiety manipulation was not salient enough in the context of reward-based learning. To assess these alternative scenarios, we performed a control behavioral experiment with new anx2 and control groups (N =13 each, see sample size estimation in *Methods and Materials*). Participants in each group performed the two training blocks 2 and 3 (**Figure 1**), but without completing a preceding baseline exploration block. In anx2, state anxiety was induced exclusively during the first training block, as in the original experiment. We found that the HRV index was significantly reduced in anx2 relative to controls during the manipulation phase (*P*_FDR_ < 0.05, Δ = 0.72, CI = [0.62, 0.83]), but not during the final training phase (block 3, *P*_FDR_ > 0.05). STAI state subscale scores rose during the anxiety manipulation in anx2 – not in controls – relative to the initial scores (within-group effect, *P*_FDR_ < 0.05, Δ = 0.68, CI = [0.59, 0.78]).

Overall the anx2 group achieved a lower average score (and final monetary reward) than control participants (*P* = 0.0256; Δ = 0.64, CI = [0.50, 0.71]). In addition, anx2 participants achieved significantly lower scores than control participants during the first training block (*P*_FDR_ < 0.05, Δ = 0.68, CI = [0.54, 0.79] **Figure 4D**), yet not during the second training block (*P*_FDR_ > 0.05). Notably, however, the degree of cvIKI or cvKvel did not differ between groups (*P*_FDR_ < 0.05, **Figure 4E-F**). The mean performance tempo, loudness and the mean learned solution during training did not significantly differ between groups, as in the main experiment (P > 0.05). Thus, removal of the baseline exploration phase led to the anxiety manipulation impairing reward-based learning, and this effect was not associated with a change in the use of task-related variability or average performance parameters.

### Bayesian learning modeling reveals the effects of state anxiety on reward-based motor learning

To assess our hypotheses regarding the mechanisms underlying participants’ performance during reward-based learning we used three different versions of a Bayesian learning model, which were based on the hierarchical Gaussian filter for continuous input data (HGF; Mathys et al., 2011, 2014). The HGF was introduced by Mathys and colleagues (2011) to model how an agent infers a hidden state in the environment (a random variable), x_1_, as well as its rate of change over time (x_2_, environmental volatility). This corresponds to a perceptual model, which is further coupled with a response model to generate responses based on those inferred states. In the HGF, beliefs about those two hierarchically-related hidden states (x_1_, x_2_) are updated given new sensory input (scores) via prediction errors (PEs). Crucial to the HGF is the weighting of the PEs by the ratio between the uncertainty of the current level and the lower level; or the inverse ratio when using precision (inverse variance or uncertainty of a distribution). Further details are provided in **Methods and Materials.**

Different implementations of the HGF have been recently used in combination with neuroimaging data to investigate how the brain processes different types of hierarchically-related prediction errors (PEs) within the framework of predictive coding (Diaconescu et al., 2017; Weber et al., 2019). The HGF can be fitted to the behavioral data of each individual participant, thus providing dynamic estimates of uncertainty and hierarchical PEs weighted by precision (precision-weighted PE or pwPE). In predictive coding models, precision is viewed as crucial for representing uncertainty and updating the posterior expectations about the hidden states (Sedley et al., 2016). In the HGF, time-varying pwPEs reflect how participants learn stimulus-outcome or response-outcome associations and their changes over time (Mathys et al., 2011, 2014; Diaconescu et al., 2017).

Here we adapted the HGF to model participants’ estimation of quantity x_1_, which represented their beliefs about the value of the *performance measure* that was rewarded (a measure of timing, keystroke velocity or a combination of both). This model also estimated participant’s beliefs about environmental volatility, x_2_. Volatility in our paradigm emerged from the multiplicity of performance-to-score mappings, as different temporal patterns of the performance with identical IKI-difference values led to the same scores. The model generated belief trajectories about the external states x_1_ and x_2_, which were further used to estimate the most likely response corresponding with those beliefs.

We implemented three HGF models corresponding with participants’ decision to modify on a trial-by-trial basis a specific performance measure – thus linking it to the rewarded hidden performance. The performance measure was (1) the degree of temporal differences between consecutive IKI values (HGF1 model), (2) the degree of differences between the loudness of subsequent keystrokes (alternative HGF2 model), or (3) a combination of both previous measures, reflecting changes both in loudness and timing (alternative HGF3 model). The rationale for using these measures in the response model was that participants were informed that the target performance was related to either a specific pattern of short and long temporal intervals, a pattern of soft and loud key presses (small and large keystroke velocities) or a combination of both. We therefore expected that participants would link the differences in IKI or Kvel (or both) between consecutive key presses to the feedback scores. In each model, the feedback scores and the trial-based performance measure were used to update model parameters, and the log model-evidence was used to optimize the model fit (Diaconescu et al., 2017; Soch and Allefeld, 2018). More details on the modeling approach can be found in the **Methods and Materials** section.

Between-group comparison focused on four variables, the mean trajectories of perceptual beliefs (μ_1_, μ_2_, means of the posterior distributions for x_1_, x_2_; **Figure 5 – figure supplement 1**), and the uncertainty about those beliefs (variances σ_1_, σ_2_; note that the inverse variance is the precision, termed π_1_, π_2_, corresponding with the confidence placed on those beliefs). In addition, the parameter ζ characterising the response model was also compared between groups. Larger values of ζ penalize choosing the response that matches current expectations for the performance measure, μ_1_.

We used Random Effects Bayesian Model Selection (BMS) to assess at the group level the three models of learning (Stephan et al., 2009; code freely available from the MACS toolbox, Soch and Allefeld, 2018). BMS provided stronger evidence for the HGF1 model, as compared to the alternative HGF2 and HGF3 models. The exceedance probability of the winner model was 0.78, and the model frequency was 62% (similar values when looking within each experimental and control group).

Using the winner model in the total population, we next evaluated between-group differences in relevant model variables across trials throughout training (**Figure 5A-C**). The main result was that anx1 relative to control participants underestimated the tendency for x_1_, that is, the degree of temporal differences between successive IKIs linked to the hidden target performance (*P*_FDR_ < 0.05, Δ = 0.71, CI = [0.59, 0.86]). This indicates a tendency towards a more isochronous performance (same IKI in consecutive intervals). By contrast, the belief estimate for phasic volatility was significantly higher in anx1 than in control participants (*P*_FDR_ < 0.05, Δ = 0.72, CI = [0.63, 0.83]). The uncertainty about environmental volatility was smaller in anx1 relative to control participants (*P*_FDR_ < 0.05, Δ = 0.67, CI = [0.52, 0.83]). Because smaller uncertainty normally leads to smaller learning rates in the HGF update equations (and smaller precision-weights on the PEs), this last outcome supports that the anx1 group did not adequately update their estimates of environmental volatility. No differences between anx2 and control participants in the estimates for x_1_ or x_2_ or their uncertainties were found. In addition, the response model parameter ζ was significantly larger in anx1 than in control participants (0.034 [0.005] and 0.026 [0.006], respectively; *P* = 0.043, Δ = 0.62, CI = [0.51, 0.82]; no differences between anx2 and control groups). Participants in the anx1 group were therefore less likely to choose the response that matched their current expectations for μ_1_ (smaller posterior probability for a response y = μ1).

**Figure 5.**
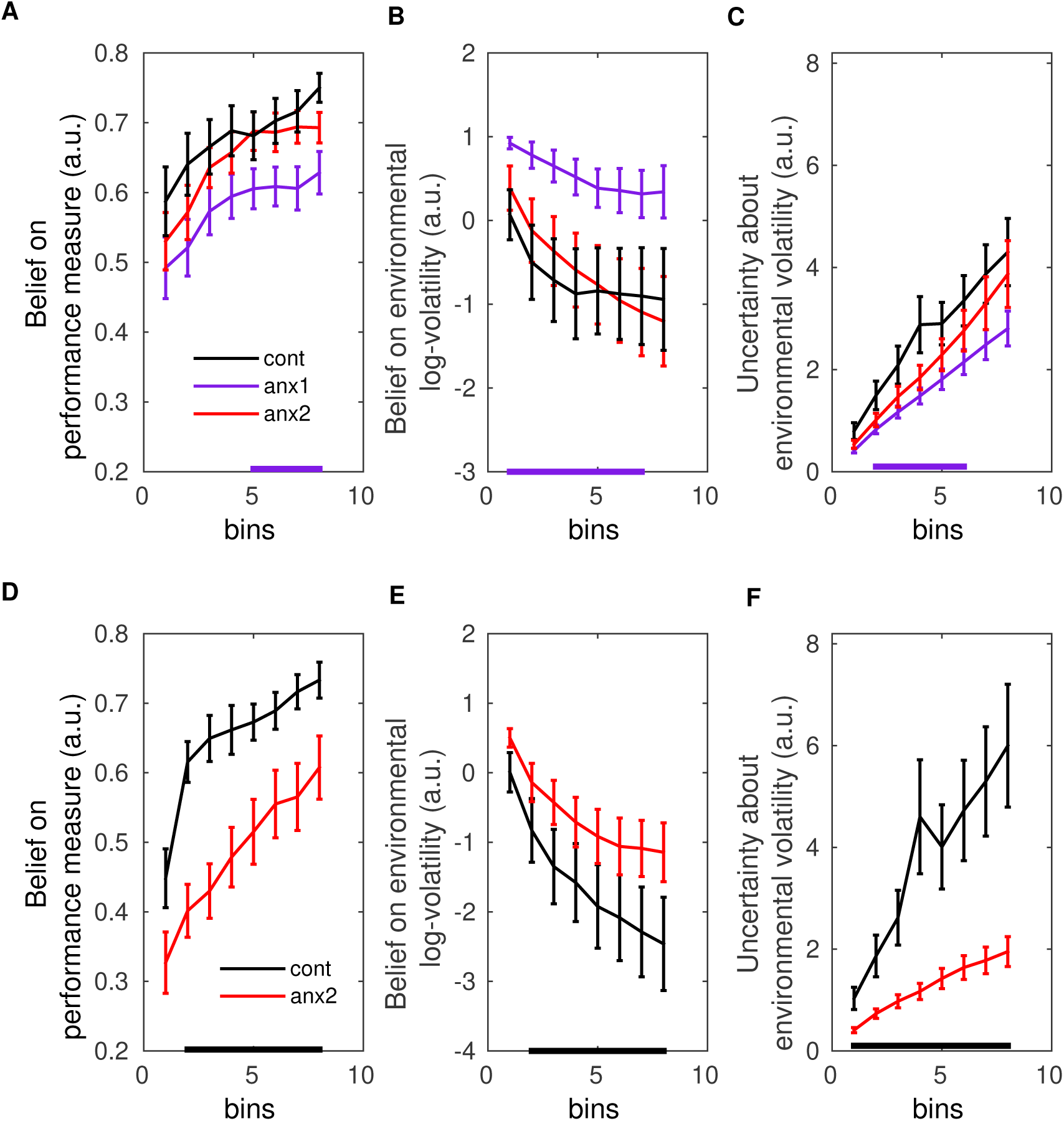
Computational modeling analysis. **(A)** In the main experiment, anx1 participants underestimated the tendency for *X_1_* (meaning their belief estimate for the degree of temporal differences between IKIs in the target performance was lower; P_FDR_ < 0.05, Δ = 0.71, CI = [0.59, 0.86], purple bar at the bottom). **(B)** By contrast, the belief estimate for environmental (phasic) volatility (*μ_2_)* was significantly higher in anx1 than control participants (P_FDR_ < 0.05, Δ = 0.72, CI = [0.63, 0.83]). **(C)** The uncertainty about environmental volatility was lower in anx1 relative to control participants (P_FDR_ < 0.05, Δ = 0.67, CI = [0.52, 0.83]), which led to smaller updates of the estimate *μ_2_.* **(D-F)** Same as (A-C) but in the separate control experiment. **(D)** The belief estimate for *x,* was lower in anx2 participants relative to control participants (P_FDR_ < 0.05, Δ = 0.75, CI = [0.67, 0.83], black bar at the botttom). **(E)** Same as (B), showing that anx2 participants overestimated the degree of environmental volatility (P_FDR_ < 0.05, Δ = 0.64, CI = [0.55, 0.73]). **(F)** Anx2 were less uncertain about their phasic volatility estimates relative to control participants (P_FDR_ < 0.05, Δ = 0.71, CI = [0.45, 0.91]). Thus, the anxiety manipulation in the control experiment biased participants to assign higher precision to their (overestimated) degree of phasic volatility.

In the second, control experiment, in which anx2 participants demonstrated a pronounced drop in scores relative to controls during the anxiety manipulation, we found that the winner model on the group level was also the HGF1 model (exceedance probability of 0.86, and model frequency of 62%). Between-group comparisons in relevant model parameters demonstrated that, similarly to anx1 participants in the main study, anx2 participants in this control experiment underestimated the tendency for x_1_ (*P*_FDR_ < 0.05, Δ = 0.75, CI = [0.67, 0.83]; **Figure 5D-F**), and overestimated the degree of phasic volatility (*P*_FDR_ < 0.05, Δ = 0.64, CI = [0.55, 0.73]). In addition, the anxiety manipulation led participants to have lower uncertainty about their phasic volatility estimates relative to control participants (*P*_FDR_ < 0.05, Δ = 0.72, CI = [0.45, 0.91]). No differences in the uncertainty about estimates for x_1_ were found. The response model parameter ζ did not differ between groups, either.

### Electrophysiological Analysis

#### State anxiety prolongs beta bursts and enhances beta power during baseline exploration

The results in **Figure 4** establish that state anxiety at baseline reduced task-related motor variability, but also subsequently led to impaired reward-based learning. We therefore sought to assess whether the anxiety-related reduced use of motor variability at baseline was associated with altered dynamics in beta-band oscillatory activity at specific time intervals during trial performance. But before investigating the dynamics of beta oscillations over time, we first looked at general averaged properties of beta activity throughout the baseline phase and their modulation by anxiety. The first measure we used was the standard averaged normalized power spectral density (PSD) of beta oscillations. Normalization of the raw PSD into decibels (dB) was carried out using as reference the average PSD from the initial rest recordings (3 min). This analysis revealed a significantly higher beta-band power in a small contralateral sensorimotor region in anx1 relative to control participants at baseline (*P* < 0.025, two-sided cluster-based permutation test, FWE-corrected. **Figure 6A-B**). In anx2 participants, the beta power in this phase was not significantly different than in controls (**Figure 6C**, *P* > 0.05). No significant between-group changes in PSD were found in lower (<13Hz) or higher (>30Hz) frequency ranges (*P* > 0.05).

**Figure 6.**
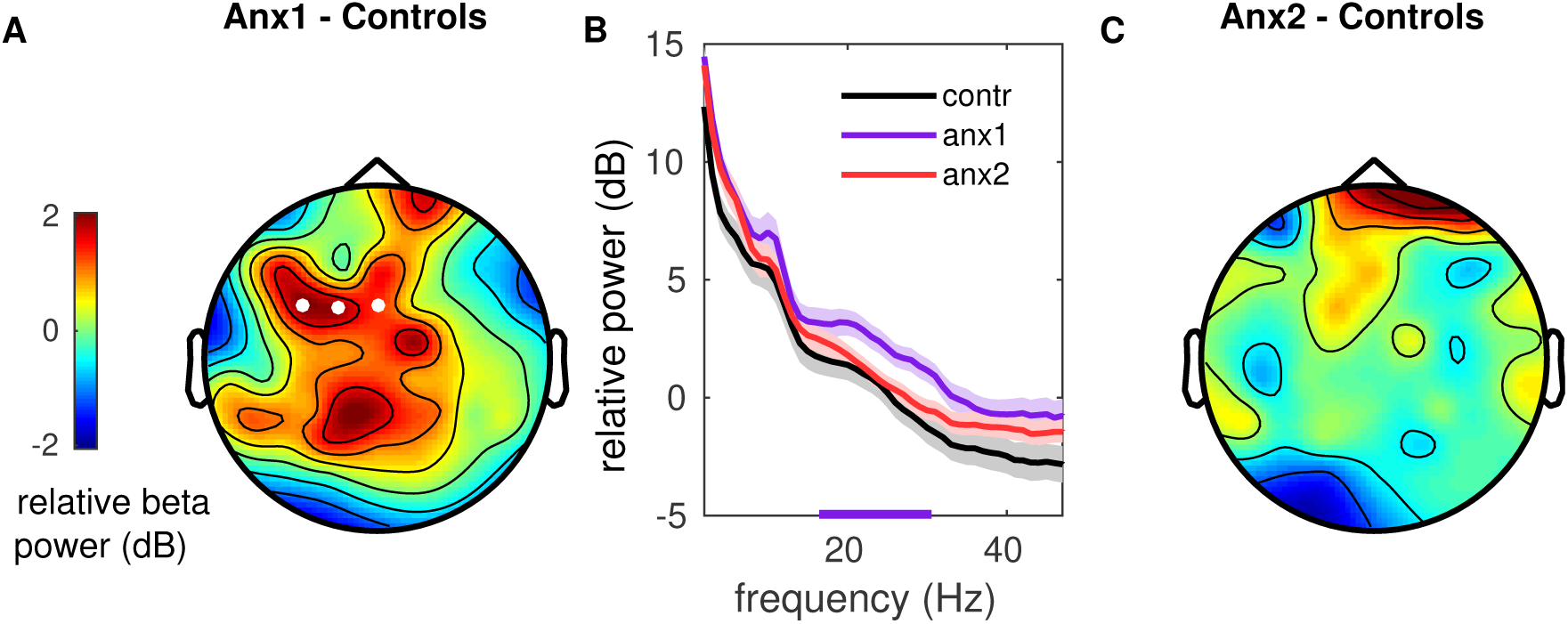
Sensorimotor beta power is modulated by anxiety during baseline exploration. **(A)** Topographical representation of the between-group difference (anxi-controls) in normalized beta-band power spectral density (PSD) in dB. A larger beta-band PSD increase was found in anx1 relative to control participants in a small cluster of contralateral sensorimotor electrodes (white dots indicate significant electrodes, two-tailed cluster-based permutation test, *P_FWE_* < 0.025). **(B)** The normalized PSD is shown within 4-45Hz for each experimental and control group after averaging within the cluster shown in (A). The purple bottom line denotes the frequency range of the significant cluster shown in (A). No significant between-group differences outside the beta range (4-i2 Hz or 3i-45 Hz) were found (*P* > 0.05). Anx2 and control participants did not differ in power modulations. Shaded areas denote mean ±SEM. **(C)** Same as (A) but for differences in beta-band PSD between anx2 and control participants. No significant clusters were found.

Next, we analyzed the between-group differences in the distribution of beta bursts extracted from the amplitude envelope of beta oscillations during baseline exploration (**Figure 7A**). This analysis was motivated by evidence from recent studies supporting that differences in the duration, rate and onset of beta bursts could account for the association between beta power and movement in humans (Little et al., 2017; Torrecillos et al., 2018). To identify burst events and assess the distribution of their duration, we applied an above-threshold detection method, which was adapted from previously described procedures (Poil et al., 2008; Tinkhauser et al., 2014; **Figure 7B**). Bursts extending for at least one cycle were selected. Using a double-logarithmic representation of the probability distribution of burst durations, we obtained a power law and extracted the slope, τ, also termed “life-time” exponent (Poil et al., 2008). Modeling work has revealed that a power law in the burst-duration distribution, reflecting that the oscillation bursts have no characteristic scale, indicates that the underlying neural dynamics operate in a state close to criticality, and thus are beneficial for information processing (Poil et al., 2008; Chialvo, 2010).

**Figure 7.**
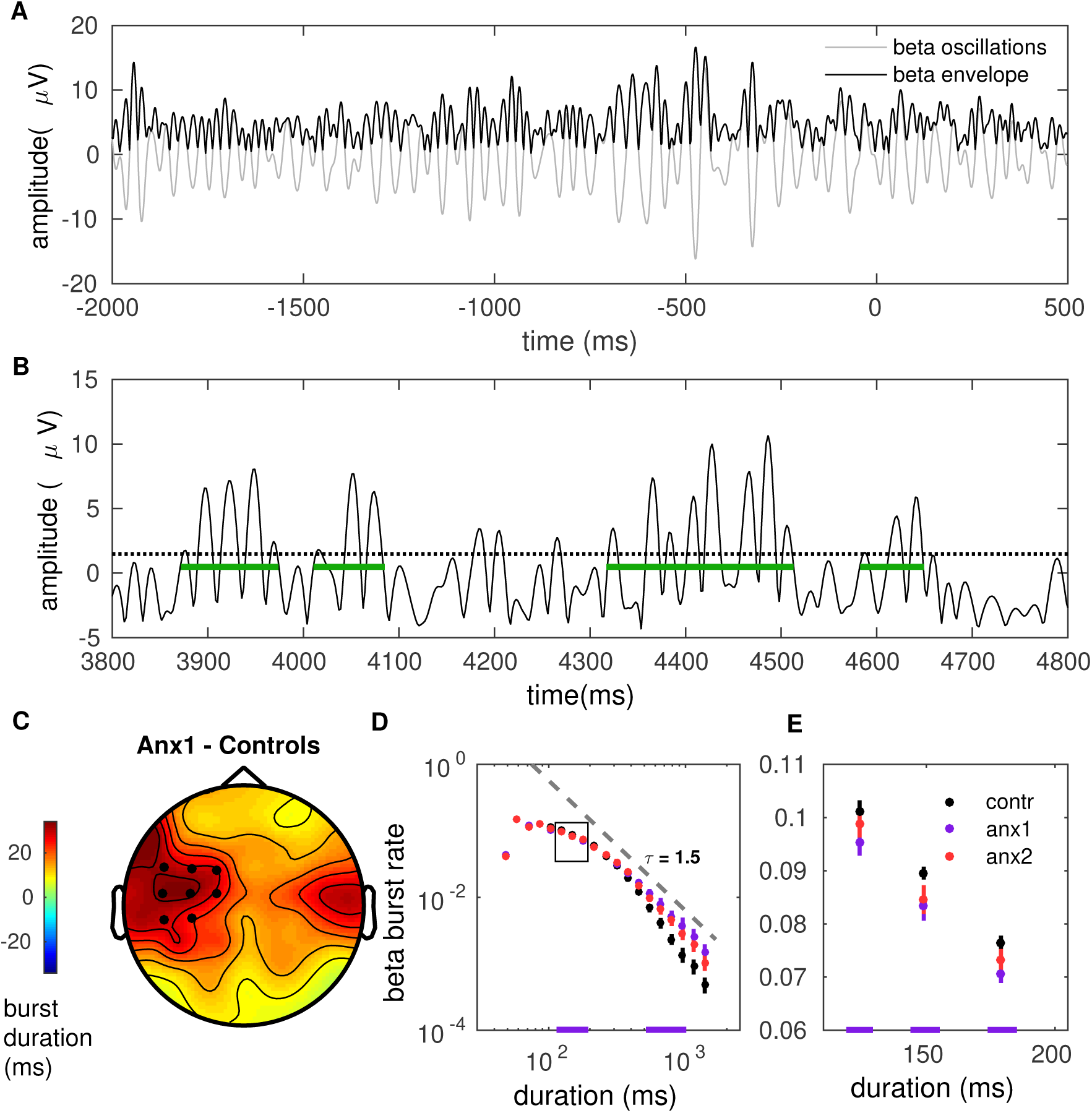
Anxiety during baseline exploration prolongs the life-time of sensorimotor beta-band oscillation bursts. **(A)** Illustration of the amplitude of beta oscillations (gray line) and amplitude envelope (black line) for one representative subject and channel. **(B)** Schematic overview of the threshold-crossing procedure to detect beta oscillation bursts. A threshold of 75% of the beta-band amplitude envelope was selected and beta bursts extending for at least one cycle were accepted. Windows of above-threshold oscillation bursts detected in the beta-band amplitude envelope (black line) are denoted by the green lines. **(C)** Scalp topography for between-group changes in the mean burst duration during baseline exploration. A significant positive cluster was found in an extended cluster of left sensorimotor electrodes, due to a longer average burst duration in anx1 than in control participants (20-30ms longer; Black dots indicate significant electrodes, two-tailed cluster-based permutation test, P_FWE_ < 0.025). **(D)** Probability distribution of beta-band oscillation-burst life-times within range 50-2000ms for each group during baseline exploration. The double-logarithmic representation reveals a power law within the fitted range (first duration bin excluded from the fit, as in Poil et al., 2008). For each power law we extracted the slope, τ, also termed life-time exponent. The dashed line illustrates a power law with τ = 1.5. Significant differences between anx1 and control participants in oscillation-burst durations are denoted by the purple line at the bottom (*P*_FDR_ < 0.05, Δ = 0.92, CI = [0.86, 0.98] for long bursts; Δ = 0.70, CI = [0.56, 0.84] for brief bursts). The rectangle highlights the area enlarged and displayed in the right panel (E). Data shown as mean and ± SEM. **(E)** Enlarged display of the region of between-group significant differences highlighted by the rectangle in (D).

In all our participants the double-logarithmic representation of the distribution of burst duration followed a decaying power-law with slope values τ in the range 1.4-1.9. Beta bursts lasted significantly longer in a contralateral sensorimotor region in anx1 as compared to control participants (**Figure 7C**, P < 0.025, FWE-corrected). The mean burst duration in these electrodes was 147 (2) ms in control participants and 168 (10) ms in the anx1 group. A further between-group comparison focusing on the distribution of burst duration demonstrated that shorter bursts were more frequent in control relative to anx1 participants (130-194ms, *P*_FDR_ < 0.05, Δ = 0.70, CI = [0.56, 0.84]; **Figure 7D-E**). By contrast, long bursts of 630-1130ms were more frequent in anx1 than control participants (*P*_FDR_ < 0.05, Δ = 0.92, CI = [0.86, 0.98]). The life-time exponents were smaller in anx1 than in the control group at left sensorimotor electrodes, corresponding with a long-tailed distribution (1.43 [0.30]; 1.70 [0.15]; *P*_FDR_ < 0.05, Δ = 0.81, CI = [0.75, 0.87]). No differences in bursts properties were found between anx2 and control participants.

We next turned to our main goal and asked whether there were between-group differences in the beta oscillatory properties at specific periods throughout the baseline exploration trials, above and beyond the general block-averaged changes reported above. This was addressed by analyzing the time course of the beta power and the beta burst rate during trial performance. Beta bursts of shorter (< 300 ms) and longer (> 500 ms) duration were assessed separately, which was motivated by previous studies linking longer beta bursts to detrimental performance (e.g. beta bursts longer than 500 ms in the basal ganglia of Parkinson’s disease patients are associated with worse motor symptoms; Tinkhauser et al., 2017). In anx1 participants the mean beta power increased after completion of the sequence performance and further following the STOP signal, and these changes were significantly more pronounced than in control participants (*P*_FDR_ < 0.05, Δ = 0.72, CI = [0.63, 0.80]; **Figure 8A**). This significant effect was localized to contralateral sensorimotor and right prefrontal channels. The rate of long oscillation bursts displayed a similar time course and topography to those of the power analysis, with an increased burst rate after movement termination and after the STOP signal in anx1 relative to control participants (*P*_FDR_ < 0.05, Δ = 0.69, CI = [0.61, 0.78]; **Figure 8B**). By contrast, brief burst events were less frequent in anx1 than in control participants, albeit exclusively during performance (*P*_FDR_ < 0.05, Δ = 0.74, CI = [0.65, 0.82]; **Figure 8C**). No significant effects were found when comparing any of these measures between anx2 to control participants.

**Figure 8.**
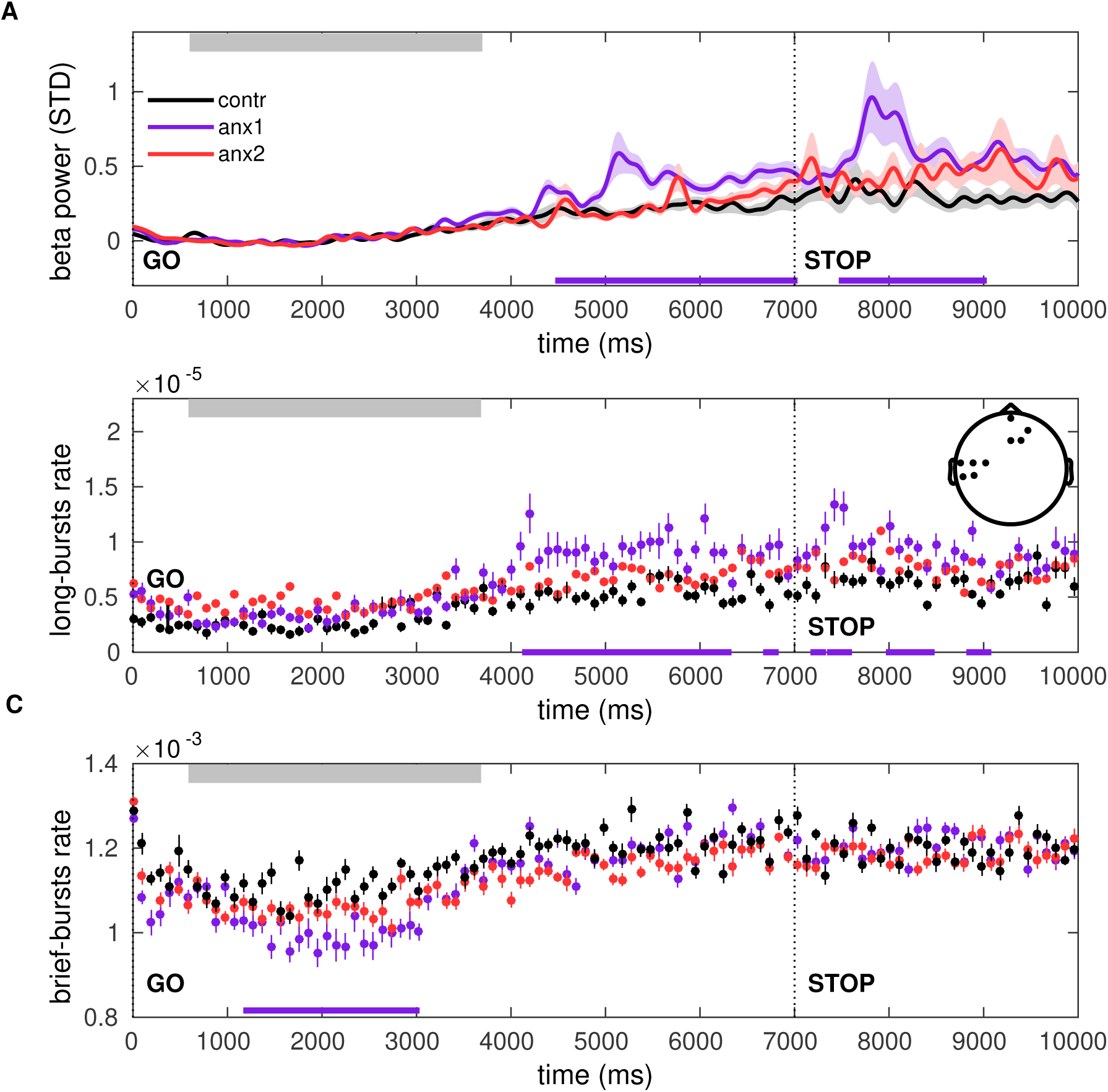
Time course of the beta power and burst rate during trials of baseline exploration. **(A)** The time representation of the beta power throughout trial performance shows two distinct time windows of increased power in participants affected by the anxiety manipulation: following sequence performance and, additionally, after the STOP signal (*P*_FDR_ < 0.05, Δ = 0.72, CI = [0.63, 0.80]; black bars at the bottom indicate the windows of significant differences). Average across sensorimotor and prefrontal electrode regions as shown in the inset in (B; *P*_FWE_ < 0.025). Shaded areas indicate the SEM around the mean. Performance of sequence1 was completed on average between 680 (50) and 3680 (i00) ms, denoted by the gray rectangle at the top. The STOP signal was displayed at 7000 ms after the GO signal, and the trial ended at 9000 ms. **(B)** The rate of oscillation bursts of longer duration (> 500 ms) exhibited a similar temporal pattern, with increased burst rate in anx1 participants following movement and the STOP signal (*P*_FDR_ < 0.05, Δ = 0.69, CI = [0.61, 0.78]). **(C)** In contrast to the rate of long bursts, the rate of brief oscillation bursts was reduced in anx1 relative to control participants, albeit during performance (*P*_FDR_ < 0.05, Δ = 0.74, CI = [0.65, 0.82]).

Additional control analyses were carried out to dissociate the separate effect of anxiety and motor variability on the time course of the beta-band oscillation properties during baseline exploration. These analyses demonstrated that, when controlling for changes in motor variability, anxiety alone could explain the findings of larger post-movement beta-band PSD and rate of longer bursts, while also explaining the reduced rate of brief bursts during performance (**Figure 8 – figure supplement 1**). Motor variability did also partially modulate the beta burst rate and power measures, after excluding anxious participants. This effect, however, had a moderate effect size and was limited to the interval after the STOP signal and contralateral sensorimotor electrodes (**Figure 8 – figure supplement 2**).

#### Reduced presence of long beta bursts promotes the update of beliefs about the volatility of motor predictions: Modulation by anxiety

During training, the general level of PSD did not differ between groups (*P*_FDR_ > 0.05; **Figure 9 – figure supplement 1A-C**), but beta-band oscillation bursts were indeed discriminative of the different experimental and control groups. Long bursts continued to be more frequent (brief bursts were less frequent) in anx1 relative to control participants in sensorimotor and prefrontal electrodes, despite the anxiety manipulation having finished in this group (**Figure 9 – figure supplement 1D-E**; *P*_FDR_ < 0.05, Δ = 0.75, CI = [0.65, 0.86]; anx1 had also smaller scaling exponents, 1.6 [0.3], than control participants, 1.9 [0.2]; *P*_FDR_ < 0.05, Δ = 0.73, CI = [0.62, 0.84]). Compared to the control group, anx2 participants exhibited a burst distribution with a longer tail, albeit exclusively in prefrontal electrodes (smaller scaling exponents in anx2, 1.69 [0.20]; *P*_FDR_ < 0.05, Δ = 0.71, CI = [0.55, 0.87]; **Figure 9 – figure supplement 2**). The mean burst duration in these prefrontal electrodes was also larger in anx2 participants (158 [20] ms in anx2, 150 [20] ms in controls, *P*_FDR_ < 0.05, Δ = 0.69, CI = [0.56, 0.82]). The lack of beta burst effects in sensorimotor electrode regions in anx2 could explain the lack of behavioral effects in this group when compared to controls.

Although **Figure 4** had established that there were no between-group differences in motor variability during training blocks (or other motor output variables), we assessed whether alterations in the beta-band measures over time during trial performance could explain the drop in scores in anx1 participants. In this group, the mean beta power increased towards the end of the sequence performance more prominently than in control participants, and this effect was significant in sensorimotor and prefrontal channels (*P*_FDR_ < 0.05, Δ = 0.67, CI = [0.56, 0.78]; **Figure 9A**). A significant increase with similar topography and latency was observed in the anx2 group relative to control participants (*P*_FDR_ < 0.05, Δ = 0.61, CI = [0.56, 0.67]). A additional and particularly pronounced enhancement in beta power appeared in anx1 and anx2 participants within 400 – 1600 ms following presentation of the feedback score. This post-feedback beta increase was significantly larger in anx1 than in the control group (*P*_FDR_ < 0.05, Δ = 0.65, CI = [0.55, 0.75]; no significant effect in anx2, *P* > 0.05).

**Figure 9.**
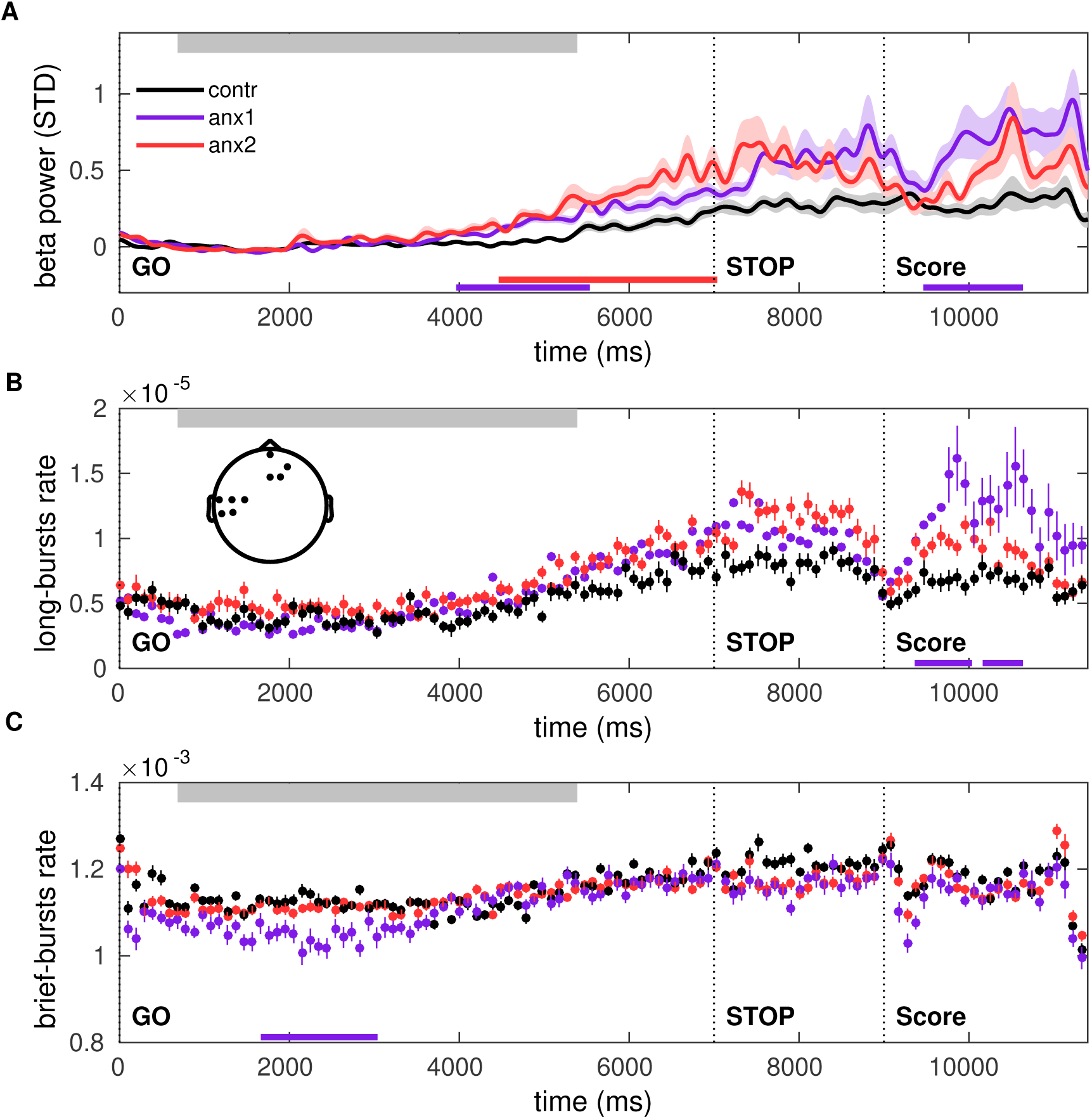
Time course of the beta power and burst rate throughout trial performance and following reward feedback. **(A)** Time course of the feedback-locked beta power during sequence performance in the training blocks, separately in anx1, anx2 and control groups. Average across sensorimotor and prefrontal electrode regions as in (B). Shaded areas indicate the SEM around the mean. Participants completed sequence2 on average between 720(30) and 5350 (100) ms, denoted by the top gray box. The STOP signal was displayed 7000 ms after the GO signal, and was followed at 9000 ms by the feedback score. This representation shows two distinct time windows of significant differences in beta activity between anx1 and control groups: at the end of the sequence performance and subsequently following feedback presentation (*P*_FDR_ < 0.05, Δ = 0.67, CI = [0.56, 0.78]; and Δ = 0.65, CI = [0.55, 0.75], respectively, denoted by the purple bar at the bottom). Anx2 participants also exhibited an enhanced beta power towards the end of the sequence performance (*P*_FDR_ < 0.05, Δ = 0.61, CI = [0.56, 0.67]). **(B)** Time course of the rate of longer (> 500 ms) oscillation bursts during sequence performance in the training blocks. Anx1 participants exhibited a prominent rise in the burst rate 400 – 1600 ms following the feedback score, which was significantly larger than the rate in control participants (*P*_FDR_ < 0.05, Δ = 0.82, CI = [0.70, 0.91]). Data display the mean and ± SEM. The topographic map indicates the electrodes of significant effects for panels (A-C; *P*_FWE_ < 0.025). **(C)** Same as (B) but showing the rate of shorter beta bursts (< 300 ms) during sequence performance in the training blocks. Between-group comparisons demonstrated a significant drop in the rate of brief oscillation bursts in anx1 participants relative to control participants at the beginning of the performance (*P*_FDR_ < 0.05, Δ = 0.70, CI = [0.56, 0.84]), yet not after the presentation of the feedback score.

Further, we found that the time course of the beta burst rate exhibited a significant increase in anx1 relative to control participants within 400 – 1600 ms following feedback presentation, similar to the power results (**Figure 9B**; *P*_FDR_ < 0.05, Δ = 0.82, CI = [0.70, 0.91]). The rate of brief oscillation bursts was, by contrast, smaller in anx1 than in control participants, albeit exclusively during performance and not during feedback processing (**Figure 9C**; *P*_FDR_ < 0.05, Δ = 0.70, CI = [0.56, 0.84]). The significant effects in anx1 participants were observed in left sensorimotor and right prefrontal electrodes, similar to the general burst effects reported in the previous section. There were no significant differences between anx2 and control groups in the rate of brief or long bursts throughout the trial (*P* > 0.05).

Having established that anx1 relative to control participants exhibited a phasic increase in beta activity and in the rate of long bursts 400 – 1600 ms following feedback presentation, we next investigated whether these post-feedback beta changes could account for the biased belief and volatility estimates in the anx1 group (**Figure 5**). In the proposed predictive coding framework, superficial pyramidal cells encode PEs weighted by precision (precision-weighed PEs or pwPEs), and these are also the signals that are thought to dominate the EEG (Friston and Kiebel, 2009). A dissociation between high (gamma > 30 Hz) and low (beta) frequency of oscillations has been proposed to correspond with the encoding of bottom-up PEs and top-down predictions, respectively (Arnal and Giraud, 2012). Operationally, however, beta oscillations have been associated with the *change* in predictions (Δμ_1_) rather than with predictions themselves (Sedley et al, 2016). In the HGF the update equations for **μ_1_** and μ_2_ are detemined exclusively by the pwPE term in that level, such that the change in predictions, Δμ_1_, is equal to pwPE (see **Methods and Materials**). Accordingly, we assessed whether the trialwise feedback-locked beta power or burst rate represented the magnitude of pwPEs in that trial that serve to update belief estimates about the performance measure (μ_1_) or the environmental volatility (μ_2_).

In each participant, we did a three-way split on the single-trial pwPE values for level 1 (termed ε_1_) and level 2 (ε_2_) and analyzed their effect on the corresponding feedback-locked beta power as a function of the participant group. This analysis focused on the interval 400-i600 ms following the feedback presentation. **Figure 10** shows as a general tendency that larger post-feedback beta activity was associated with smaller pwPEs. A 2 x 3 non-parametric factorial analysis with factors Group (anx1, controls) and Magnitude of ε_1_ (small, medium, large) revealed a significant main effect of Group, as expected (*P* = 0.01; factorial analysis with synchronized rearrangements, Basso et al., 2007; **Figure 10A**). No significant main effect of Magnitude or interaction effect was found (*P* > 0.05). A similar analysis carried out for ε_2_ indicated that the main effects of Group and Magnitude of ε_2_ were significant (*P* = 0.01 and 0.045, respectively). Thus, in addition to the post-feedback beta power being modulated by the group factor, the results supported that the increase in beta activity following feedback presentation represented the magnitude of the precision-weighted PEs that drive updates about volatility estimates; and independently of the group factor.

**Figure 10.**
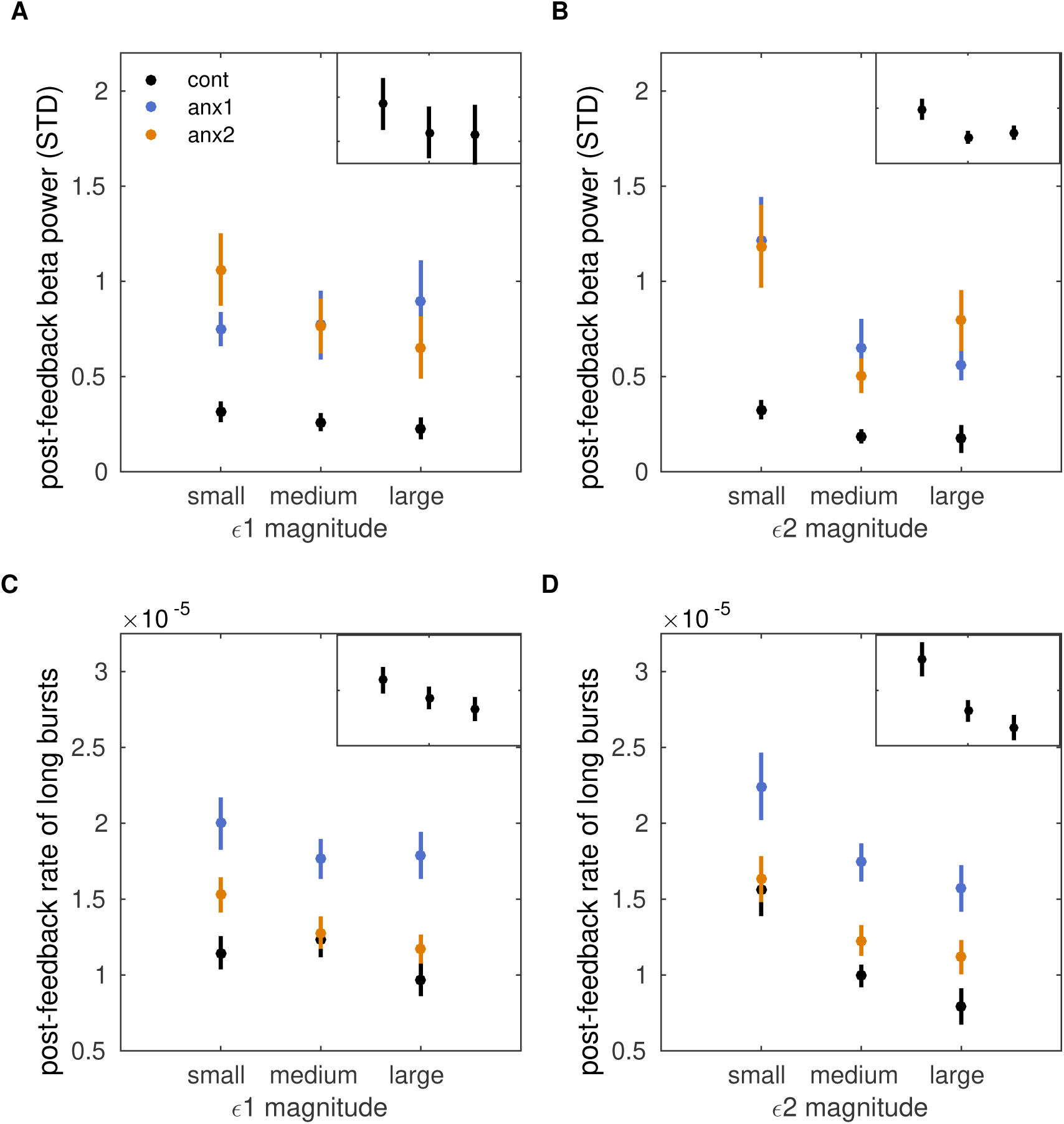
Post-feedback increases in the rate of long oscillation burst represent attenuated precision-weighted prediction errors about volatility estimates. **(A)** Average beta power within 400-1600 ms following feedback presentation in training blocks, sorted by the magnitude of precision-weighted PEs (pwPEs) for level 1 (ε_1_). Although the post-feedback beta power in control participants decreased with increasing magnitude of ε_1_, and so did the grand-average across all 60 participants (top right inset), our non-parametric 3 x 2 factorial analysis did not reveal significant main effects for factors Magnitude of ε_1_ or Group (anx1, controls); neither did we find interaction effects (see main text). No effects when using anx2 in the Group factor either. **(B)** Same as (A) but for the magnitude of pwPEs for level 2 (ε2), driving belief updates about volatility. No significant main effects or interactions were found. **(C)** Grand-average of the rate of post-feedback long beta bursts sorted by the magnitude of trial-wise pwPEs driving belief estimates about the performance measure. A 2 x 3 non-parametric factorial analysis with factors Group (anx1, controls) and Magnitude of ε_1_ revealed a significant main effect of Group (*P* = 0.028). A trend of significance was found for factor Magnitude (*P* = 0.065). **(D)** Both main effects were significant when considering the pwPEs of the second level, ε_2_ (*P* = 0.032 and 0.027). This result links the higher post-feedback rate of long-lived oscillation bursts in anx1 with reduced updates about volatility.

The analysis of the rate of long oscillation bursts revealed a pattern consistent with the beta power results, with smaller pwPEs being associated with a larger burst rate. A 2 x 3 non-parametric factorial analysis with factors Group (anx1, controls) and Magnitude of ε_1_ revealed a significant main effect of Group (*P* = 0.028; **Figure 10C**). A trend of significance was found for factor Magnitude (*P* = 0.065). Both main effects were significant when considering the pwPEs of the second level, ε_2_ (*P* = 0.032 and 0.027, for Group and Magnitude factors, respectively; **Figure 10D**). The results highlight that the more frequent presence of long-lived beta bursts following feedback, as found in anx1 (**Figure 10C-D**), could be linked to a reduced update in predictions about volatility estimates and, to a lesser degree (trend), estimates about the performance measure. The rate of brief oscillation bursts following the outcome presentation was not modulated by pwPEs (**Figure 10 – figure supplement 1**). Neither did we find an association between raw changes in (non-weighted) PEs and changes in beta burst or power properties (P > 0.05).

## Discussion

The results revealed several interrelated mechanisms through which state anxiety impairs reward-based motor learning. First, state anxiety induced biases about the hidden performance goal and it’s stability throughout time. Second, anxiety led to an underestimation of uncertainty about volatility, thereby attenuating the update of beliefs about this quantity. In addition, we found that state anxiety reduced motor variability at baseline, decreasing performance in the subsequent reward-based learning phase. On the neural level, bursts sensorimotor beta oscillations, a marker of physiological beta (Feingold et al., 2015), lasted longer under the effect of anxiety during baseline exploration, resembling recent findings of abnormal burst duration in movement disorders (Tinkhauser et al., 2017). The anxiety-induced higher rate of long burst events at baseline extended to prefrontal electrodes and also to the following training phase, where additional phasic trial-by-trial feedback-locked increases in this measure accounted for the biases in the update of beliefs about volatility. These results provide the first evidence for state anxiety inducing changes in the distribution of sensorimotor and prefrontal beta bursts, thereby leading to deficits in the update of beliefs about volatility during reward-based motor learning.

Evidence from our main experiment supported that the finding of anxiety-related reduced motor variability at baseline was associated with the outcome of subsequently impaired learning from reward. These results validate previous accounts on the relationship between motor variability and Bayesian inference (Wu et al. 2014). In addition, the association between larger baseline task-related variability and higher scores during the following training phase extends results on the faciliatory effect of exploration on motor learning, at least in tasks that require learning from reinforcement (Wu et al., 2014; Pekny et al., 2015; Dhawale et al., 2017; see also critical view in He et al., 2017).

Crucially, however, the lack of between-group differences in the use of task-related variability during training in both experiments indicates that this measure could not account for the anxiety-related deficits in reward-based learning. In fact, the evidence from the control experiment supported that state anxiety can impair learning from reward by directly influencing computations of uncertainty and belief estimates independently of changes in prior or concurrent variability. Our Bayesian learning model revealed that what impaired participants subjected to the anxiety manipulation in both experiments from achieving high scores was an underestimation of the target performance measure, as well as an overestimation of environmental volatiliy, which led them to estimate the hidden goal as being more unstable throughout time. In addition, they had smaller uncertainty about environmental volatility. This implies that they considered their estimation of volatility to be more precise, and requiring smaller updates (the update equations are directly proportional to the uncertainty estimate at that level). The results align well with recent computational work in decision-making tasks, showing that high trait anxiety leads to deficits in uncertainty estimates and adaptation to the changing statistical properties in the environment (Browning et al, 2015; Huang et al., 2017). Our findings thus provide the first evidence that computational mechanisms similar to those described for trait anxiety and decision-making underlie the effect of temporary anxious states on motor learning. This might be particularly the case in the context of learning from rewards, such as feedback about success or failure, which is considered one of the fundamental processes through which motor learning is accomplished (Wolpert et al., 2011).

Previous studies manipulating psychological stress and anxiety to assess motor learning showed both a deleterious and faciliatory effect (Hordacre et al., 2016; Vine et al., 2013; Bellomo et al., 2018). Differences in experimental tasks, which often assess motor learning during or after high-stress situations but not during anxiety induction in anticipation of a stressor, could account for the previous mixed results. Here, we adhered to the neurobiological definition of anxiety as a psychological and physiological response to an upcoming diffuse and unpredictable threat (Bishop, 2007; Grupe and Nitschke, 2013). Accordingly, anxiety was induced using the threat of an upcoming public speaking task (Feldman et al., 2004; Lang et al., 2015), and was associated with a drop in the HRV and an increase in state anxiety scores during the targeted blocks. Although the average state anxiety scores were not particularly high, they were significantly higher during the targeted phases than during the initial resting state phase. Future studies should however use more impactful stressors to study the effect of the full spectrum of state (and trait) anxiety on motor learning (Bellomo et al., 2018).

What is the relationship between the expression of motor variability and state anxiety? As hypothesized, state anxiety at baseline reduced the use of variability across trials. This converges with recent evidence demonstrating that anxiety leads to ritualistic behavior (repetition, redundancy, rigidity of movements) to regain a sense of control (Lang et al., 2015). The outcome also aligns well with animal studies where evidence shows a reduction in motor exploration when stakes are high (high-reward situations, social context; Dhawale et al., 2017; Kao et al., 2008; Woolley et al., 2014). These interpretations, however, seem to stand in contrast with our findings in anx2 participants, which were affected by the anxiety manipulation during training yet this had no effect on their use of motor variability or achieved scores when compared to controls. The control experiment clarified this issue by demonstrating that removal of a baseline motor exploration phase leads to anxiety diminishing reward-based learning through changes in belief and volatility estimates and deficits in processing uncertainty – and independently of changes in concurrent motor variability. Thus, the evidence combined supports that the normal use of baseline variability in anx2 participants in the main experiment might have protected them from the effects of the anxiety manipulation, favouring the interpretation that initial unconstrained exploration is important for *subsequent* successful motor learning.

Some considerations should be taken into account. Task-related motor variability might be pivotal for learning from reinforcement or reward signals (Sutton and Barto, 1998; Wu et al., 2014; Dhawale et al., 2017), whereas in other contexts, such as during motor adaptation, the evidence is conflicting (He et al., 2017, Shin et al., 2016). An additional consideration is that higher levels of motor variability could reflect both an intentional pursuit of an explorative regime; or, an unintentional higher level of motor noise, similarly to previous work (Wu et al., 2014; Pekny et al., 2015). A recent study established that motor learning is improved by the use of intended exploration, not motor noise (Chen et al., 2017). Our paradigm cannot dissociate between intended and unintended exploration. This limitation will be addressed in future work by using a separate baseline phase with regular performance to assess motor noise as a measure of unintended exploration. Lastly, we used a reward-based motor learning paradigm in which different performances could provide the same feedback score. Thus, a high expression of task-related motor variability during training could lead the participants to perceive the task as volatile. The rationale for using this task was to explore the effect of state anxiety on volatility estimates, as recent work demonstrates that anxiety primarily affects learning in volatile conditions (Browning et al., 2015; Huang et al., 2017). Volatility in our task, however, could be detrimental for learning regardless of the participant group, which the correlation results across all 60 participants confirmed. Further analyses revealed that the mean learned performance and the degree of motor variability during training were not different between groups, supporting that these are not confounding factors that could explain the reward-based-learning group results. Instead, our findings underscore that computational mechanisms related to belief and uncertainty estimates are the main factors driving the effects of concurrent or prior state anxiety on reward-based motor learning.

On the neural level, an important finding was that anxiety at baseline increased the power of beta oscillations, the duration of beta bursts and the rate of long beta bursts (long-tailed distribution). The increases in power and rate of long-lived bursts manifested after completion of the sequence, reflecting an anxiety-related enhancement of the post-movement beta rebound (Kilavik et al., 2012, 2013). This effect was observed in a region of contralateral sensorimotor and right prefrontal channels, and could be explained by anxiety alone, despite a small effect of motor variability on the modulation of these neural changes across sensorimotor electrodes. Our analyses did not provide a detailed anatomical localization of the effect, yet the findings in sensorimotor regions partly contributing to changes in motor variability are consistent with the involvement of premotor and motor cortex in driving motor variability and learning, as previously reported in animal studies (Churchland et al., 2006; Mandelblat-Cerf et al, 2009; Santos et al., 2015). The results also converge with the representation in the premotor cortex of temporal and sequential aspects of rhythmic performance (Crowe et al., 2014; Kornysheva and Diedrichsen, 2014).

During training, the concurrent anxiety manipulation in anx2 participants affected the HRV and increased the presence of long bursts exclusively in prefrontal electrodes. This outcome aligns with the finding of prefrontal involvement in the emergence and maintainance of anxiety states (Davidson, 2002; Bishop et al., 2007; Grupe and Nitschke, 2013). Here we showed that manipulating state anxiety shifts the physiological distribution of beta bursts in prefrontal regions towards a long-tailed distribution, characterized by more frequent long bursts. In addition, the lack of beta burst effects in sensorimotor regions in this group is in agreement with the absence of behavioral effects when compared with control participants. An unexpected result was the mainteinance in anx1 of higher rates of long bursts across sensorimotor and prefrontal electrodes during training, which extended from the previous phase. Thus, our results revealed that in the context of motor learning anxious states can induce changes in sensorimotor beta power and burst distribution, which are maintained after physiological measures of anxiety return to baseline, thus continuing to affect relevant behavioral parameters. Anxiety has been shown to modulate different oscillatory bands depending on the context, such as gamma activity in visual areas and amygdala during processing fearful faces (Schneider et al., 2018), alpha activity in response to processing emotional faces (Knyazev et al., 2008) or theta activity during rumination (Andersen et al., 2009). Beta-band oscillations could be particularly relevant to flesh out the effects of anxiety on performance during motor tasks.

Mechanistically, phasic trial-by-trial feedback-locked changes in the burst distribution were related to the computational biases in belief updates and uncertainty estimates found in anx1 participants, and explained their poorer performance during reward-based learning. Specifically, a higher rate of long beta bursts and increased power following feedback in this group were associated with a reduced update in predictions about volatility estimates. The post-feedback increase in the long burst rate also showed a tendency (trend) to represent updates in predictions about the performance measure. The computational quantity that determines the update of predictions in our Bayesian model is the precision-weighted PEs, which here were inversely related to the rate of long beta bursts and beta power. Raw changes in (non-weighted) PEs were not associated with changes in beta burst or power properties. This outcome is in line with the predictive coding hypothesis that PEs are mediated by gamma oscillations, whereas neuronal signalling of predictions is mediated by lower frequencies (e.g., alpha 8-12Hz, Friston and Litvak, 2015). Further studies point to beta oscillations as the relevant cortical oscillatory rhythm associated with encoding predictions, although the evidence so far is scarce (Arnal and Giraud, 2012). More recently, beta oscillations have been associated with the *change* to predictions rather than with predictions themselves (Sedley et al, 2016), which is consistent our findings. In line with these results, a post-performance increase in beta power during motor adaptation is considered to index confidence in priors and thus a reduced tendency to change the ongoing motor comand (Tan et al., 2014). More generally, beta oscillations along cortico-basal ganglia networks have been proposed to gate incoming information to modulate behavior (Leventhal et al., 2012) and to maintain the current motor state (Engel and Fries, 2010). Consequently, the phasic increase in beta power and the rate of beta bursts following feedback presentation could represent neural states facilitating encoding of pwPEs and update in predictions about relevant quantities.

Our findings support that assessing neural activity in sensorimotor regions is crucial to understand the effects of anxiety on motor learning and to determine mechanisms above and beyond the role of prefrontal control of attention in mediating the effects of anxiety on cognitive and perceptual tasks (Bishop et al., 2007; 2009; Eyseneck, 2007). Our data imply that the combination of Bayesian learning models and analysis of oscillation burst properties can help better understand the mechanisms through which anxiety modulates motor learning. Future studies should investigate how the brain circuits involved in anxiety interact with motor regions to affect motor learning. In addition, assessing burst properties across both beta and gamma frequency ranges would further allow us to delineate and dissociate the neural mechanisms responsible for anxiety biasing decision-making and motor learning.

## Methods and Materials

### Participants and sample size estimation

Sixty right-handed healthy volunteers (37 females) aged 18 to 44 (mean 27 years, SEM, 1) participated in the main study. In a second, control experiment, 26 right-handed healthy participants (16 females, mean age: 25.8, SEM 1, range 19-40) took part in the study. Participants gave written informed consent prior to the start of the experiment, which had been approved by the local Ethics Committee at Goldsmiths University. Participants received a base rate of either course credits or money (£15; equally distributed across groups) and were able to earn an additional sum up to £20 during the task depending on their performance.

We used pilot data from a behavioral study using the same motor task to estimate the minimum sample sizes for a statistical power of 0.95, with an α of 0.05, using the MATLAB (The MathWorks, Inc., MA, USA) function sampsizepwr. In the pilot study we had one control and one experimental group of 20 participants each. In the experimental group we manipulated the reward structure during the first reward-based learning block (in this block feedback scores did not count towards the final average monetary reward). For each behavioral measure (motor variability and mean score), we extracted the standard deviation (sd) of the joint distribution from both groups and the mean value of each separate distribution (e.g. m1: control, m2: experimental), which provided the following minimum sample sizes:

Between-group comparison of behavioral parameters (using 2-tailed t-test): MinSamplSizeA = sampsizepwr(‘t’,[m1 sd],m2, 0.95) = 18-20 participants.

Accordingly, we recruited 20 participants for each group in the main experiment. Next, using the behavioral data from the anxiety and control groups in the current main experiment, we estimated the minimum sample size for the second, behavioral control experiment:

Between-group comparison of behavioral parameters (using 2-tailed t-test): MinSamplSizeA = sampsizepwr(‘t’,[m1 sd],m2, 0.95) = 13 participants.

Therefore for the second control experiment we recruited 13 participants in each group.

### Apparatus

Participants were seated at a digital piano (Yamaha Digital Piano P-255, London, United Kingdom) and in front of a PC monitor in a light-dimmed room. They sat comfortably in an arm-chair with their forearms resting on the armrests of the chair. The screen displayed the instructions, feedback and visual cues for start and end of a trial (**Figure 1A**). Participants were asked to place four fingers of their right hand (excluding the thumb) comfortably on 4 pre-defined keys on the keyboard. Performance information was transmitted and saved as Musical Instrument Digital Interface (MIDI) data, which provided time onsets of keystrokes relative to the previous one (inter-keystroke-interval – IKI in ms), MIDI velocities (related to the loudness, in arbitrary units, a.u.), and MIDI note numbers that corresponded to the pitch. The experiment was run using Visual Basic and additional parallel port and MIDI libraries.

### Materials and Experimental design

In all blocks, participants initiated the trial by pressing a pre-defined key with their left index finger. After a jittered interval of 1-2 s, a green ellipse appeared in the centre of the screen representing the GO signal for task execution (**Figure 1**). Participants had 7 s to perform the sequence which was ample time to complete it before the green circle turned red indicating the end of the execution time. If participants failed to perform the sequence in the correct order or initiated the sequence before the GO signal, the screen turned yellow. In blocks 2 and 3 during training, performance-based feedback in form of a score between 0 and 100 was displayed on the screen 2 s after the red ellipse, that is, 9 s from the beginning of the trial. The scores provided participants with information regarding the target performance.

The performance measure that was rewarded during training was the Euclidean norm of the vector corresponding with the pattern of temporal differences between adjacent IKIs for a trial-specific performance. Here we denote the vector norm by ||**Δz**||, with **Δz** being the vector of differences, **Δz** = (*z_2_* – *z_1_, z*_3_ – *z*_2_, …, *z*_n_ – *z*_n−1_), and *z_1_* representing the IKI at each keystroke (*i* = 1, 2.., n). Note that IKI values themselves represent the difference between the onset of consecutive keystrokes, and therefore **Δz** indicates a vector of *differences of differences.* Specifically, the target value of the performance measure was a vector norm of 1.9596 (e.g. one of the maximally rewarded performances leading to this vector norm of IKI-differences would consist of IKI values: [0.2, 1, 0.2, 1, 0,2, 1, 0.2] s; that is a combinaiton of short and long intervals). The score was computed in each trial using a measure of proximity between the *target* vector norm **||Δz^t^**|| and the norm of the *performed* pattern of IKI differences ||**Δz^p^**||, using the following expression:

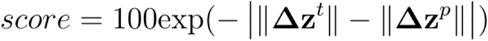

In practice, different temporal patterns leading to the same vector norm ||**Δz^p^**|| could achieve the same score. Participants were unaware of the existence of various solutions. Higher exploration across trials during training could thus reveal that several IKI patterns were similarly rewarded. To account for this possibility, the perceived rate of change of the hidden goal (environmental volatility) during training was estimated and incorporated into our mathematical description of the relationship between performance and reward (see below).

### Anxiety Manipulation

Anxiety was induced during block1 performance in group anx1, and during block2 performance in the anx2 group by informing participants about the need to give a 2-minute speech to a panel of experts about an unknown art object at the end of that block (Lang et al., 2015). We specified that they would first see the object at the end of the block (it was a copy of Wassily Kandinsky’ Reciprocal Accords [1942]) and would have 2 min to prepare for the presentation. Participants were told that the panel of experts would take notes during their speech and would be standing in front of the testing room (due to the EEG setup participants had to remain seated in front of the piano). Following the 2-min preparation period, participants were informed that due to the momentary absence of panel members they instead had to present in front of the lab members. Participants in the control group had the task to describe the artistic object to themselves, not in front of a panel of experts. They were informed about this secondary task before the beginning of block1.

### Assessment of State Anxiety

To assess state anxiety we acquired two types of data: (1) the short version of the Spielberger State-Trait Anxiety Inventory (STAI, state scale X1, 20 items; Spielberger, 1970) and (2) a continuous electrocardiogram (ECG, see EEG, ECG and MIDI recording session). The STAI X1 subscale was presented four times throughout the experiment. A baseline assessment at the start of the experiment before the resting state recording was followed by an assessment immediately before each experimental block to determine changes in anxiety levels. In addition, a continuous ECG recording was obtained during the resting state and three experimental blocks to assess changes in autonomic nervous system responses. The indexes of heart rate variability (HRV, coefficient of variation of the inter-beat-interval) and mean heart rate (HR) were evaluated, as their reduction has been linked to changes in anxiety state due to a stressor (Feldman, 2004).

### Computational Model

Here we provide details on the computational Bayesian model we adopted to estimate participant-specific belief trajectories, their uncertainty and the precision-weighted PEs. The model was implemented using the HGF toolbox for MATLAB® (http://www.translationalneuromodeling.org/tapas/). The model consists of a perceptual and a response model, representing an agent (a Bayesian observer) who generates behavioral responses based on a sequence of sensory inputs (scores) it receives. As general notation, we let lower case italics denote scalars (*x),* which can be further characterised by a trial superscript *x^k^* and a subscript *i* denoting the level in the hierarchy 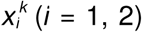. We use lower case bold font for vectors with n components, **x**.

The HGF corresponds to the perceptual model, representing a hierarchical belief updating process, i.e., a process that infers hierarchically related environmental states that give rise to sensory inputs (Stefanics, 2011; Mathys et al., 2014). It then generates belief trajectories about external states, such as the reward value of an action or a stimulus. In the version for continuous inputs we use here (see Mathys et al., 2014; function `tapas_hgf.m`), learning occurs in two hierarchically coupled levels (*x_1_, x_2_),* one for “perceptual” beliefs (*x_1_*: the rewarded performance measure), and the phasic volatility of those beliefs (*x_2_*). These two levels evolve as coupled Gaussian random walks, with the lower level coupled to the higher level through its variance (inverse precision). The Gaussian random walk at each level *X_i_* is determined by its posterior mean (*μ_i_*) and its variance (*σ_i_*). Further, the variance of the lower level, *x_1_*, depends on *x_2_* through an exponential function

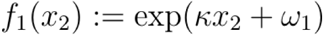

*κ* and *ω_1_* are model parameters that are estimated in each participant by fitting the HGF model to the experimental data (scores and responses) using Variational Bayes. At the top level, the variance is typically fixed to a constant parameter, *ϑ*. The specific coupling between levels indicated above has the advantage of allowing simple variational inversion of the model and the derivation of one-step update equations under a mean-field approximation. Importantly, the update equations for the posterior mean at level *i* and for trial *k* depend on the prediction errors weighted by precision (or uncertainty) according to the following expression:

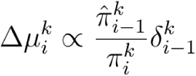

The first term in the above expression is the change in the expectation or current belief 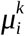 for state *X_i_,* and the previous expectation in triak *k-1* 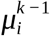. This difference term is proportional to the prediction error of the level below, 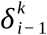, representing the difference between the expectation 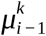 and the prediction 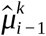 of the level below 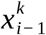. The prediction error is weighted by the ratio between the prediction of the precision of the level below, 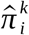, and the precision of the current belief, 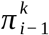. This expression illustrates that higher uncertainty in the current level (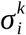, lower 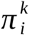 in the denominator) leads to faster update of beliefs.

Next, we selected the posterior mean *μ_1_* of the continuous variable *x_1_* representing participants’ beliefs about the value of the performance measure that was rewarded (a measure of timing, keystroke velocity or a combination of both; see below), and fed it to a separate continuous *response model* to link those estimates to participant’s responses. As response model we chose the Gaussian noise model for responses on a continuous scale (function `gaussian_obs.m` in the TAPAS toolbox). This model is defined by a Gaussian distribution centered at the difference between participants’ current estimates for *X_1_, μ_1_,* and their responses *y:*

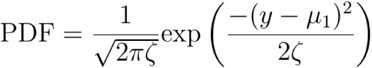

The posterior probability for a participant choosing response *y* in this model is therefore largest when the response matches the most likely value of *μ_1_* according to its current belief. This Gaussian distribution is normalized with parameter *ζ*, which penalises the choice of a specific response *y* (decreasing the posterior probability). The participant-specific estimates of parameter *ζ* were also provided by the HGF toolbox.

Each of the three implemented full (perceptual + response) models corresponded to participants’ decision to modify on a trial-by-trial basis a specific performance measure – thus linking it to the rewarded hidden performance. The performance measure was (1) the degree of temporal differences between consecutive keystrokes (HGF1 model), (2) the degree of differences between loudness of subsequent keystrokes (alternative HGF2 model), or (3) a combination of both previous measures, reflecting changes both in loudness and timing (alternative HGF3 model). The rationale for using these measures in the response model was that participants were informed that the target performance was related to either a pattern of short and long temporal intervals, small and large keystroke velocities or a combination of both. We therefore assummed that participants would link the differences in IKI or Kvel (or both) between consecutive key presses to the feedback scores. Accordingly, for model HGF1 we fed as responses the following normalized quantity (range 0-1).

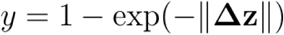

Here, ||**Δz**|| represents the norm of the vector of differences between adjacent IKI (≡ *z)* values for a trial-specific performance (See **Stimulus Materials** and score computation). Model HGF2 corresponded to participants’ decision to modify the pattern of differences in loudness (*z* ≡ Kvel) between successive keystrokes: **Δz** = (*Kvel_2_* – *Kvel_1_, Kvel_3_* – *Kvel_2_, …, Kvel_n_-Kvel_n-1_)·* MIDI velocity values within 0-127 were normalized to the range 0-1 (Kvel/127), as IKI values fell within 0-1 s. Lastly, model HGF3 implemented the scenario in which participants decided to vary both the pattern of IKIs and differences in Kvel on a trial by trial basis. In this case, the argument of the exponential was the mean between the norm of **Δz** in model HGF1 and HG2, respectively:

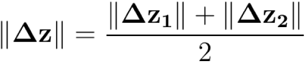

The priors on the model parameters (*ω, κ* ϑ) and on the initial expected states 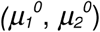 are provided in **Table 1**. All priors are Gaussian distributions in the space in which they are estimated and are therefore determined by their mean and variance. The variance is relatively broad to let the priors be modified by the series of inputs (scores). Quantities that need to be positive (e.g. variance or uncertainty of belief trajectories) are estimated in the log-space, whereas general unbounded quantities are estimated in their original space.

**Table 1.** Means and variances of the priors on perceptual parameters and initial values. Priors on the parameters and initial values of HGF perceptual model for continuous inputs were the default prior values defined in function `tapas_hgf_config.m` from the HGF toolbox. The continuous inputs here were the trial-by-trial scores that participants received, normalized to the Ο-Ι range. Quantities estimated in the logarithmic space are denoted by log(). Prior mean and variance for 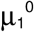, as well as the prior mean for 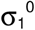 and ωι were defined by the initial input values. For the remaining quantities, the prior mean and variance were pre-defined according to the values indicated in the table. The prior means for ωι and 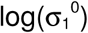) were related as the variance of variable xi in the HGF is a function of the upper level according to the expression 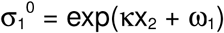.

|  | Prior mean | Prior variance |
| --- | --- | --- |
| $\log(\kappa)$ | $\log(1)$ | 0 |
| $\omega_1$ | Log-variance of the first 20 input scores (median = $\log[0.02] = -3.9$) | 16 |
| $\log(\vartheta)$ | -4 | 16 |
| $\mu_1^0$ | Value of first input score (median = 0.21 in the total population of 60 participants) | Variance of the first 20 inputs scores (median = 0.05 across all participants) |
| $\log(\sigma_1^0)$ | Log-variance of the first 20 input scores (median = $\log[0.02]$) | 1 |
| $\mu_2^0$ | 1 | 0 |
| $\log(\sigma_2^0)$ | $\log(0.1)$ | 1 |

We used Random Effects Bayesian Model Selection (BMS) to assess at the group level the three models of learning (Stephan et al., 2009; code freely available from the MACS toolbox, Soch and Allefeld, 2018). BMS provided the estimated model frequencies, that is, how frequently each model is optimal in the sample of participants and the exceedance probabilities, reflecting the posterior probability that one model is more frequent than the others (Soch et al., 2016).

### EEG, ECG and MIDI recording

EEG and ECG signals were recorded using a 64-channel (extended international 10-20 system) EEG system (ActiveTwo, BioSemi Inc.) placed in an electromagnetically shielded room. During the recording, the data were high-pass filtered at 0.16 Hz. The vertical and horizontal eye-movements (EOG) were monitored by electrodes above and below the right eye and from the outer canthi of both eyes, respectively. Additional external electrodes were placed on both left and right earlobes as reference. The ECG was recorded using two external channels with a bipolar ECG lead II configuration. The sampling frequency was 512 Hz. Onsets of visual stimuli, key presses and metronome beats were automatically documented with markers in the EEG file. The performance was additionally recorded as MIDI files using the software Visual Basic and a standard MIDI sequencer program on a Windows Computer.

### EEG and ECG pre-processing

We used MATLAB and the FieldTrip toolbox (Oostenveld et al., 2011) for visualization, filtering and independent component analysis (ICA; runica). The EEG data were highpass-filtered at 0.5 Hz (Hamming windowed sinc finite impulse response [FIR] filter, 3380 points) and notch-filtered at 50 Hz (847 points). Artifact components in the EEG data related to eye blinks, eye movements and the cardiac-field artifact were identified using ICA. Following IC inspection, we used the EEGLAB toolbox (Delorme & Makeig, 2004) to interpolate missing or noisy channels using spherical interpolation. Finally, we transformed the data into common average reference.

Analysis of the ECG data with FieldTrip focused on detection of the QRS-complex to extract the R-peak latencies of each heartbeat and use them to evaluate the HRV and HR measures in each experimental block.

### Analysis of power spectral density

We first assessed the standard power spectral density (PSD, in mV^2^/Hz) of the continuous raw data in each performance block and separately for each group. The PSD was computed with the standard fast Fourier Transform (Welch method, Hanning window of 1s with 50% overlap). The raw PSD estimation was normalized into decibels (dB) with the average PSD from the initial rest recordings (3 min). Specifically, the normalized PSD during the performance blocks was calculated as ten times the base-10 logarithm of the quotient between the performance-block PSD and the resting state power.

In addition, we assessed the time course of the spectral power over time during performance. Trials during sequence performance were extracted from −1 to 11 s locked to the GO signal. This interval included the STOP signal (red ellipse), which was displayed at 7 s, and – exclusively in training blocks – the score feedback, which was presented at 9 s. Thus, epochs were effectively also locked to the STOP and Score signals. Artefact-free EEG epochs were decomposed into their time-frequency representations using a 7-cycle Morlet wavelet in successive overlapping windows of 1 seconds within the total 12s-epoch. The frequency domain was sampled within the beta range from 13 to 30 Hz at 1 Hz intervals. The time-varying spectral power was computed as the squared norm of the complex wavelet transform, after averaging across trials within the beta range. This measure of spectral power was further averaged within the beta-band frequency bins and normalized by substracting the mean and dividing by the standard deviation of the power estimate in the pre-movement baseline period ([-1, 0] s prior to the GO signal).

### Extraction of beta-band oscillation bursts

We assessed the distribution, onset and duration of oscillation bursts in the time series of beta-band amplitude envelope. We followed a procedure adapted from previous work to identify oscillation bursts (Poil et al., 2008; Tinkhauser et al. 2017). In brief, we used as threshold the 75% percentile of the amplitude envelope of beta oscillations. Amplitude values above this threshold were considered to be part of an oscillation burst if they extended for at least one cycle (50 ms: as a compromise between the duration of one 13 Hz-cycle [76 ms] and 30 Hz-cycle [33 ms]). Threshold-crossings that were separated by less than 25 ms were considered to be part of the same oscillation burst. As an additional threshold the median amplitude was used in a control analysis, which revealed similar results, as expected from previous work (Poil et al., 2008). Importantly, because threshold crossings are affected by the signal-to-noise ratio in the recording, which could vary between the different performance blocks, we selected a common threshold from the initial rest recordings separately for each participant (Tinkhauser et al. 2017). Distributions of the rate of oscillation bursts per duration were estimated using equidistant binning on a logarithmic axis with 20 bins between 50-2000 ms.

General burst properties were assessed in baseline and training blocks separately, first as averaged values within the full block-related recording, and next as phasic changes over time during trial performance. Trial-based analysis focused on the interval 0-11000 ms following the GO signal, which included the time window following the STOP signal (at 7000 ms: baseline and training blocks) and the score feedback (at 9000 ms: training blocks).

### Statistical Analysis

Statistical analysis of behavioral and neural measures focused on the separate comparison between each experimental group and the control group (contrasts: anx1 – controls, anx2 – controls). Differences between experimental groups anx1-anx2 were evaluated exclusively concerning the overall achieved monetary reward. When appropriate, we tested main effects and interactions in factorial analyses using N x M synchronized rearragements (Basso et al., 2007). The factorial analysis was complemented with non-parametric permutation tests to assess differences between conditions or between groups in the statistical analysis of behavioral or neural measures. To evaluate differences between sets of multi-channel EEG signals corresponding to two conditions or groups, we used two-sided cluster-based permutation tests (Maris & Oostenveld, 2007) and an alpha level of 0.025. Control of the family-wise error (FWE) rate was implemented in these tests to account for the problem of multiple comparison (Maris & Oostenveld, 2007). When multiple testing was performed with permutation tests and synchronized rearrangements, we implemented a control of the false discovery rate (FDR) at level q = 0.05 using an adaptive linear step-up procedure (Benjamini et al., 2006). This control provided an adapted threshold *p-value* (termed *P*_FDR_).

Non-parametric effect size estimators were used in association with our nonparametric statistics, following Grissom and Kim (2012). In the case of between-subject comparisons, the standard probability of superiority, Δ, was used. Δ is defined as the proportion of greater values in sample B relative to A, when values in samples A and B are not paired: Δ = P (A > B). Δ ranges within 0-1. The total number of comparisons is the product of the size of sample A and sample B (Ntot = sizeA*sizeB), and therefore, Δ = N (A > B) / Ntot. In the case of ties, Δ is corrected by subtracting in the denominator the number of ties from the total number of comparisons (Ntot – Nties). For within-subject comparisons, we used the probability of superiority for dependent samples, Δ_dep_, which is the proportion of all within-subject (paired) comparisons in which the values for condition B are larger than for condition A. Confidence intervals (CI) for Δ were estimated with bootstrap methods (Ruscio & Mullen, 2012).

## Acknowledgements

This research is supported by the British Academy through grant R134610 to M.H.R. We thank Marta Garcia Huesca and Silvia Aguirre for carrying out some of EEG experiments.

## Author Contribution

Maria Herrojo Ruiz: Conceptualization, Supervision, Methodology, Data analysis, Coding, Visualisation, Funding acquisition, Project administration, Writing—original draft, review and editing.

Sebastian Sporn: Conceptualization, Data acquisition, Writing—review and editing.

Thomas P. Hein: Data acquisition, Writing—review and editing.

---

**Figure 4 – figure supplement 1.**
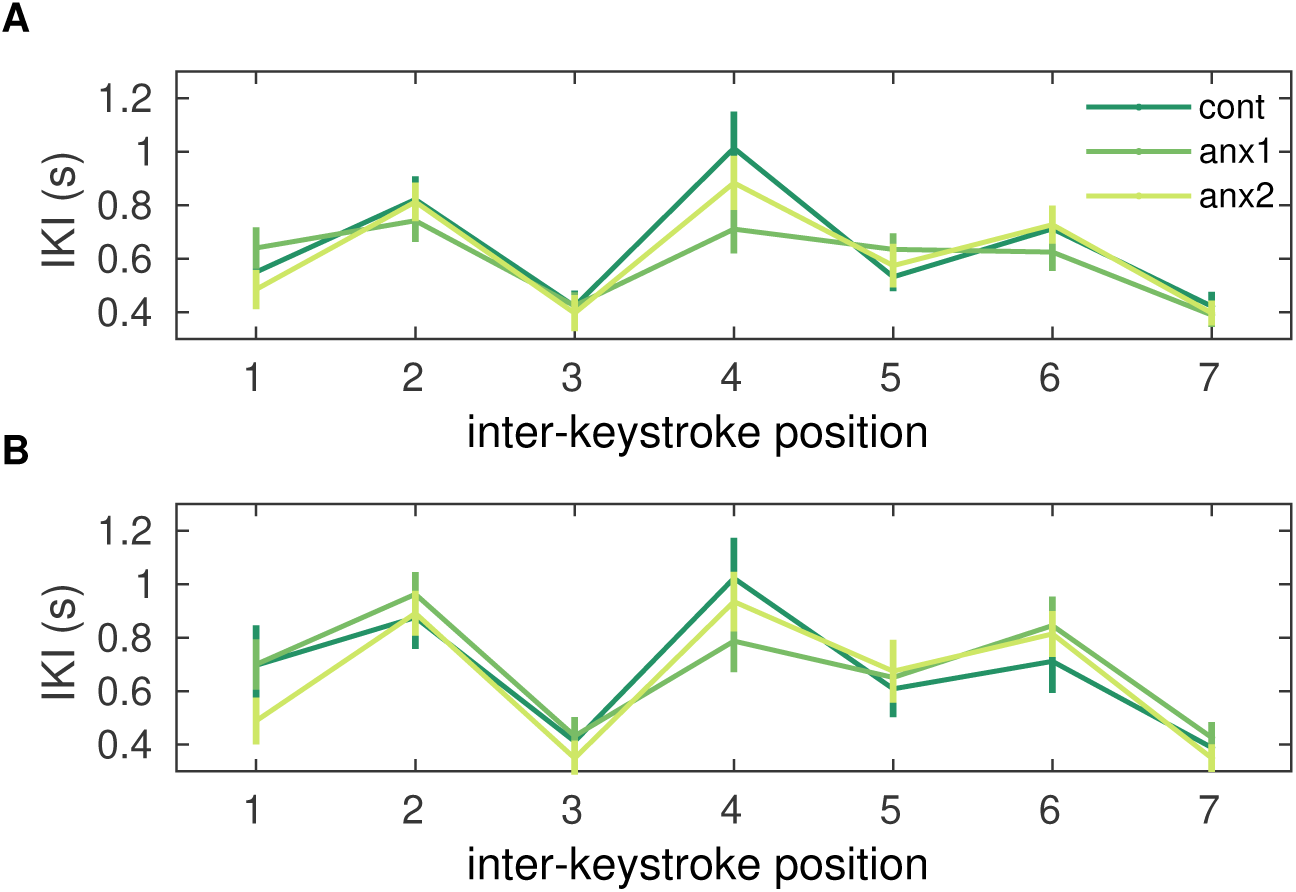
Mean learned solution in each group. On average, the learned performance in each group was not significantly different, either during the first **(A)** or second **(B)** training block (*P* > 0.05).

**Figure 5 – figure supplement 1.**
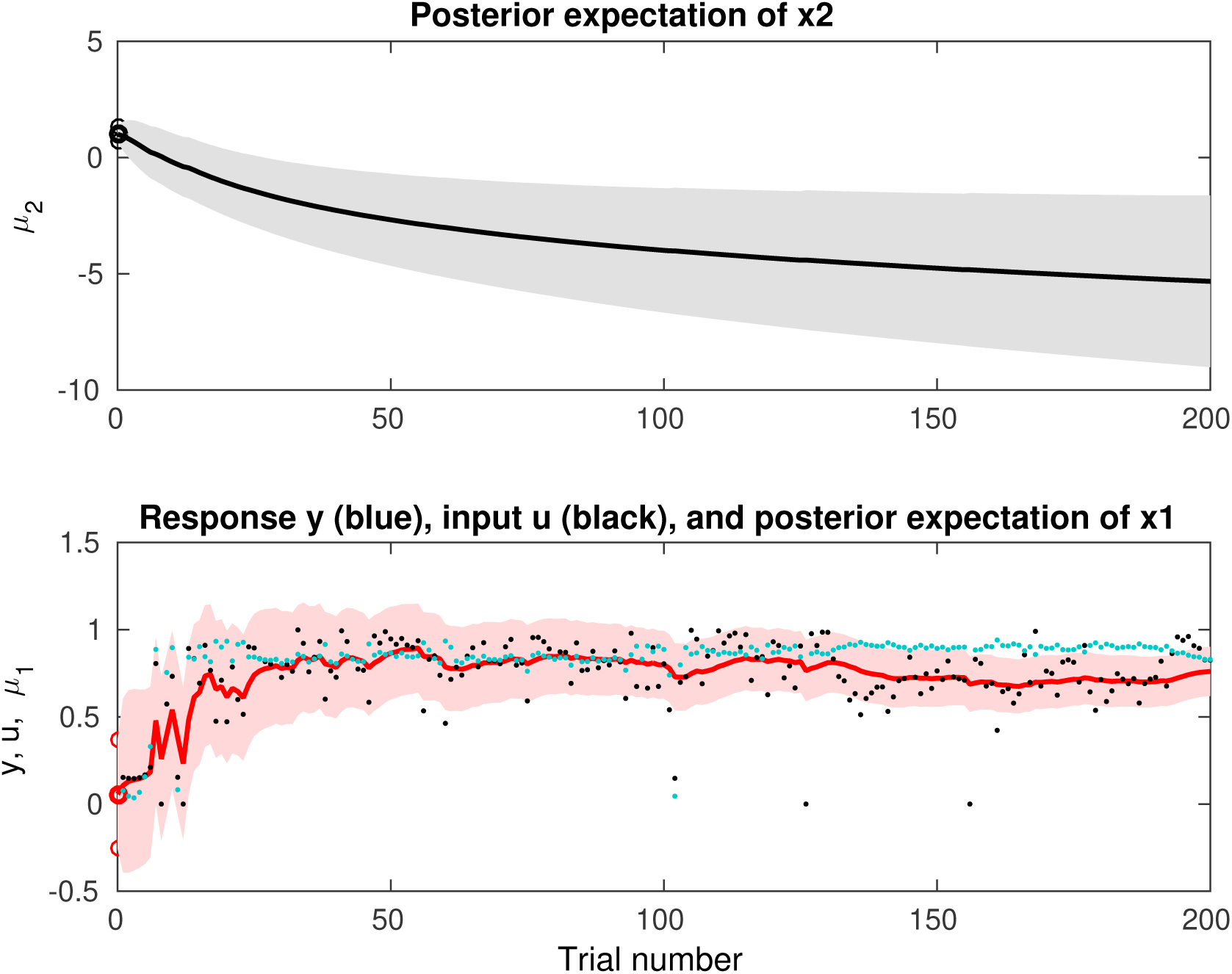
HGF Model Trajectories. Single-trial model-based estimates of the belief trajectories at the lower and higher HGF level in a representative subject. Bottom: Posterior expectation μ_1_ of the target performance measure, x_1_ Trialwise inputs (scores, denoted by the black dots) and responses (normalized degree of temporal differences between consecutive inter-keystroke-intervals; denoted by the blue dots) are shown. Top: Posterior expectation μ_2_ of the log-volatility of the environment, x_2_, representing the estimated rate of change in the lower quantity, x_1_. Shaded areas indicate the standard deviation of the probability distribution.

**Figure 8 – figure supplement 1.**
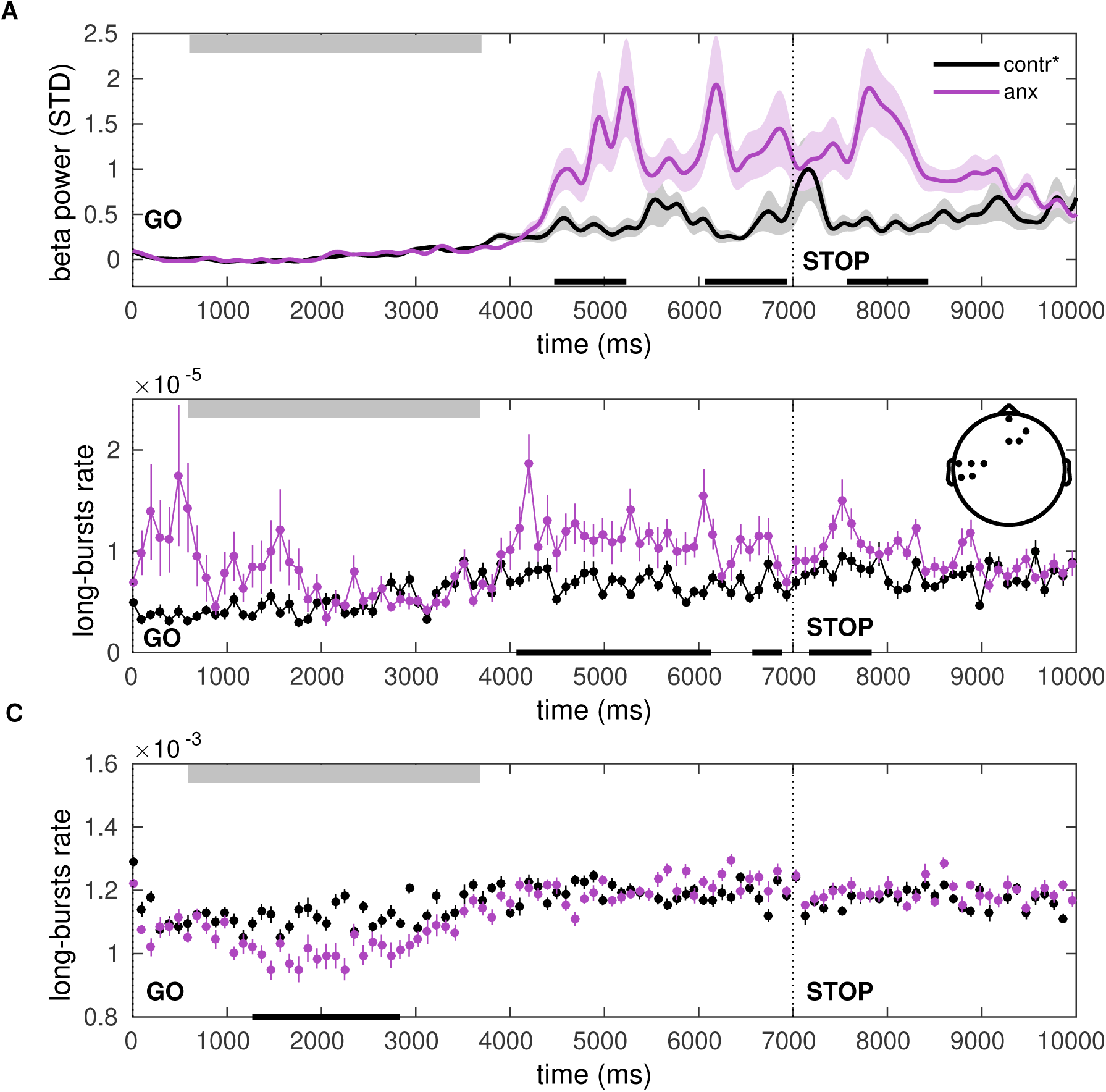
Post-movement increases in the beta-band amplitude and burst rate can be explained by state anxiety. (A-C) A separate control analysis was carried out to determine the influence of the anxiety manipulation alone on the beta PSD and burst rate properties, after controlling for changes in motor variability (cvIKI). Panels (A-C) are similar to panels (A-C) in Figure 8, but for a comparison between anx1 and participants from an extended control group (contr*, including control and anx2 participants, who were not affected by anxiety at baseline), after matching them in motor variability. Significant between-group differences are denoted by the black bar at the bottom (*P*_FDR_ < 0.05, large effect sizes, Δ = 0.81, CI = [0.72, 0.90]). This analysis revealed effects in the same windows as the primary between-group analysis shown in Figure 8.

**Figure 8 – figure supplement 2.**
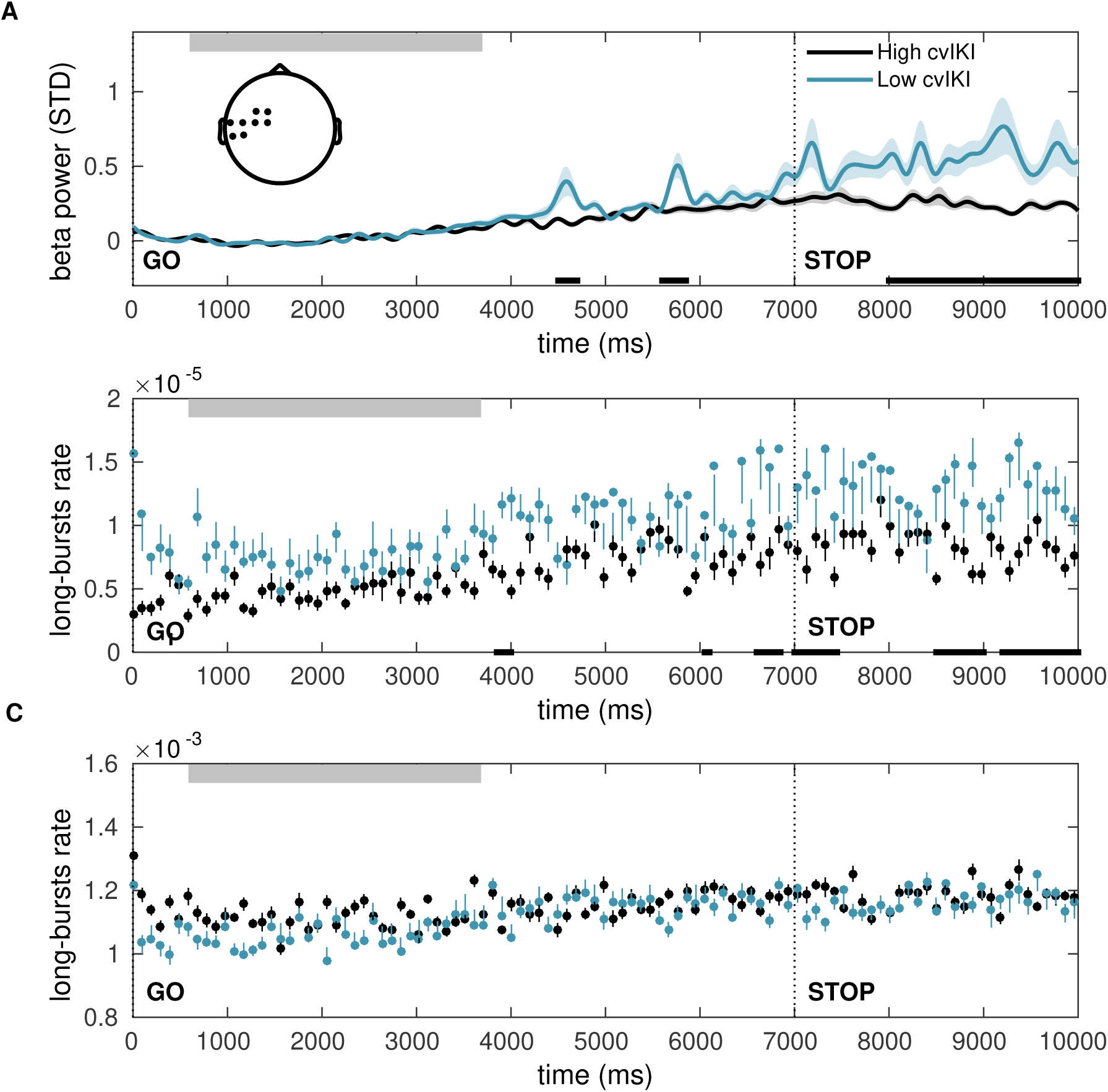
Changes in motor variability without concurrent changes in state anxiety partially account for the observed alterations in post-movement beta amplitude and burst rate. **(A-C)**. Same as **Figure 8 and Figure 8-figure supplement 1**, but in a control analysis performed to assess the effect of motor variability on beta PSD changes at baseline, independently of the anxiety manipulation. We compared participants selected from the extended control group (control + anx2) after doing a median split of the group based on their degree of temporal variability (cvIKI). Between-group differences were associated with small effect sizes (*P*_FDR_ < 0.05, Δ = 0.56, CI = [0.51, 0.62]; black bars at the bottom) and exclusively in sensorimotor electrodes (topographic map; *P*_FWE_ < 0.025).

**Figure 9 – figure supplement 1.**
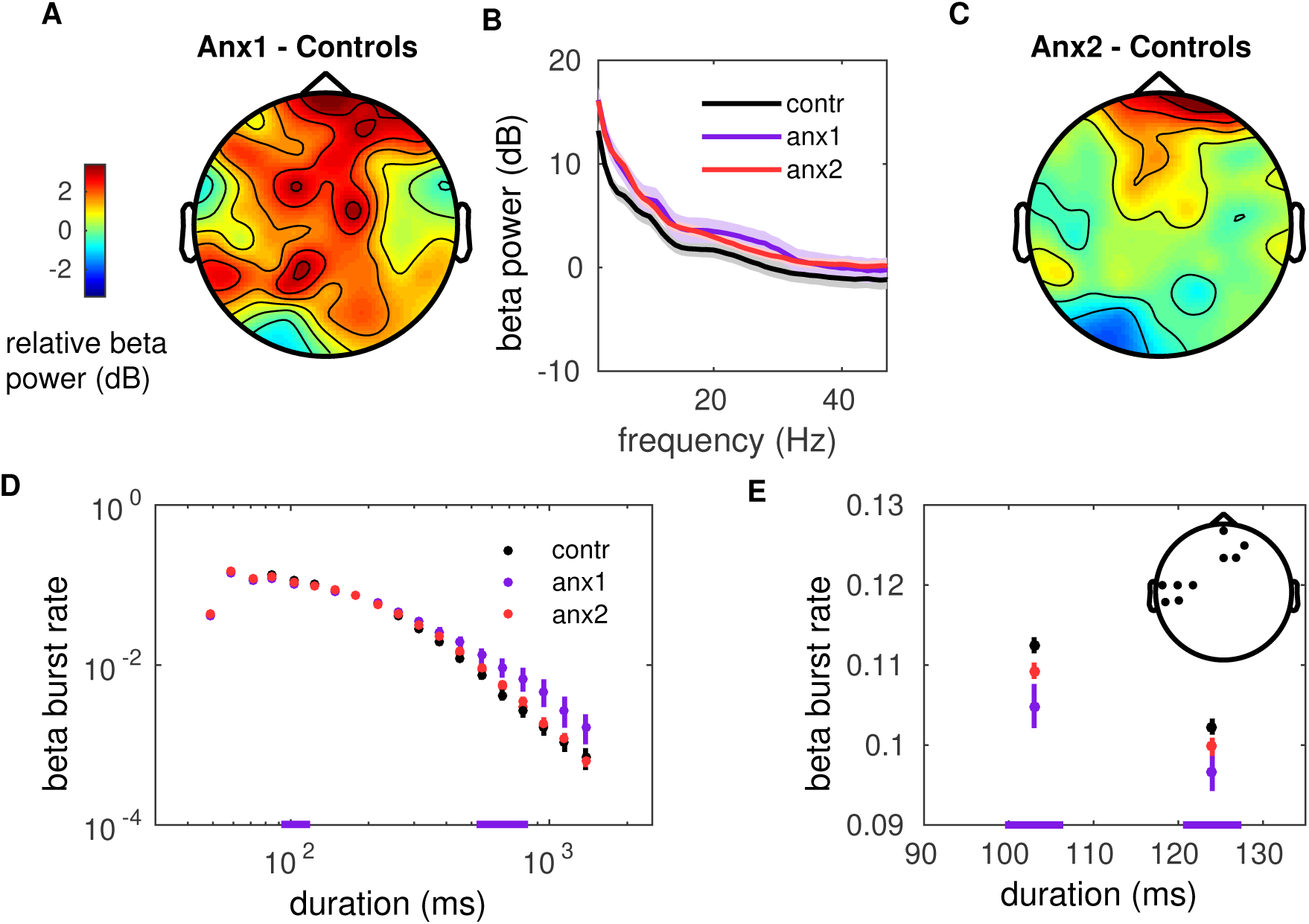
Beta power spectral density and burst rate during reward-based learning. **(A-C)**. During training, the general level of normalized PSD did not differ between groups (*P*_FDR_ > 0.05). The training-related PSD was normalized into decibels (dB) with the PSD of the initial resting state recording. **(D)** Probability distribution of beta-band oscillation-burst life-times within range 50-2000 ms for each group during training blocks. The double-logarithmic representation highlights that longer-tailed distributions were observed in anx1 participants, who exhibited more frequent long bursts and less frequent brief bursts than the control group (*P*_FDR_ < 0.05, Δ = 0.75, CI = [0.65, 0.86]; purple bars at the bottom). Data shown as mean and ± SEM. **(E)** Enlarged display of the region of significant differences for brief oscillation bursts shown in (D). The topographic map indicates the electrodes where the significant between-group rate effects were localized: left sensorimotor and right prefrontal electrode regions (*P*_FWE_ < 0.025).

**Figure 9 – figure supplement 2.**
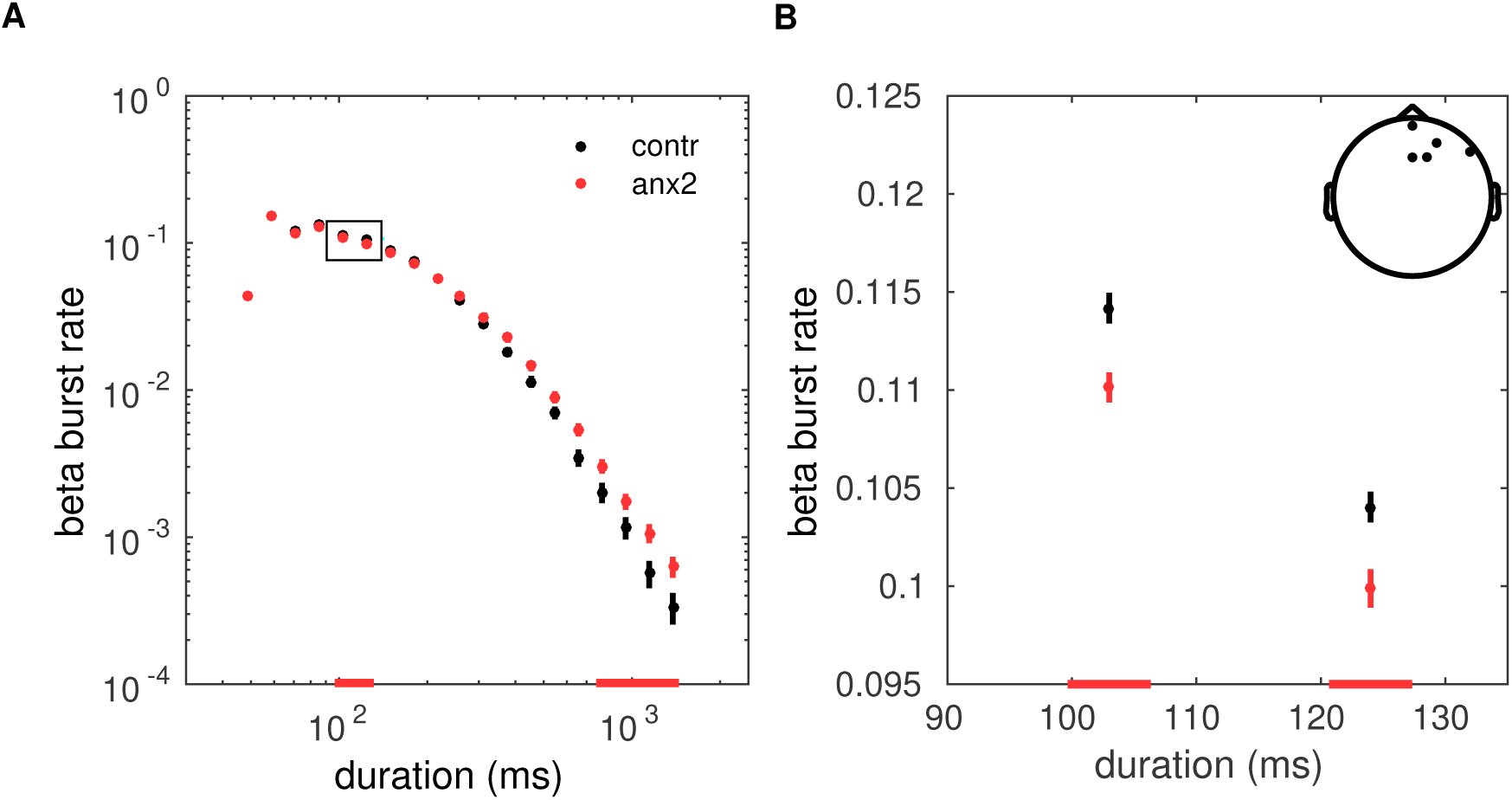
Effect of the anxiety manipulation on prefrontal burts during training. **(A-B)** Similar to Figure 9 – figure supplement 1, but for the analysis of between-group differences in anx2 and control participants. Participants in the anx2 group did not show behavioral differences as compared to the control group. Corresponding with this result, the effect of the anxiety manipulation in the anx2 group on the burst rate was limited to prefrontal electrodes, and did not extend to sensorimotor regions (*P*_FDR_ < 0.05, Δ = 0.71, CI = [0.55, 0.87]).

**Figure 10 – figure supplement 1.**
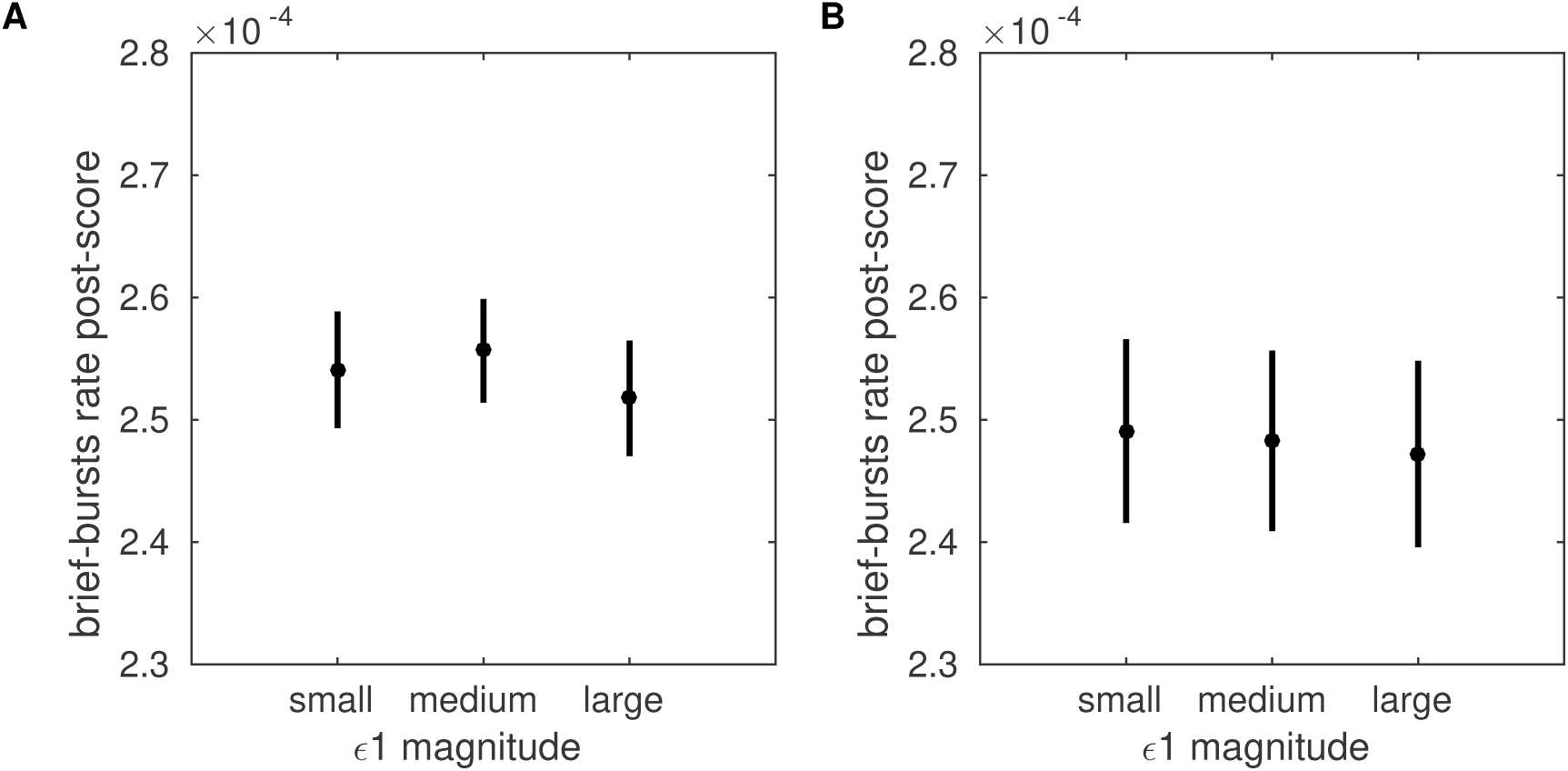
The rate of brief beta bursts following feedback is not modulated by the magnitude of precision-weighted prediction errors. Grand-average of the rate of post-feedback brief beta bursts sorted by the magnitude of trial-wise precision-weighted PEs driving belief estimates **(A)** and estimates about environmental volatility **(B)**. No significant differences were found (P > 0.05).

